# *Cola* Subgenus *Disticha* subg. nov. (Malvaceae-Sterculiaceae) of tropical Africa, a synoptic taxonomic revision with seven new species

**DOI:** 10.1101/2022.08.16.504105

**Authors:** Martin Cheek

## Abstract

A new subgenus, Subg. *Disticha* is erected for 14 species of simple-leaved *Cola* sharing unique characters within the genus which are presumed synapomorphies: distichous phyllotaxy; petioles lacking pulvini; petioles of uniformly short length, <10 mm long; male flowers with short androphores <1(– 2) mm long; stamens 4 – 5; stem indumentum where present, dominated by short simple translucent hairs (except in *C. philipi-jonesii* where stellate), and in many species dark purple to black stems with contrasting bright white lenticels. All species have fruit with small globose, unsculptured orange mericarps c. 1 cm diam. (where fruits are known). The species of this subgenus range from SE Nigeria in the west to coastal Kenya and Tanzania in the east and Malawi in the south, but are absent from the Congo basin. Species diversity is highest in the Cross-Sanaga and Eastern Arc-Coastal Forest biogeographic areas, consistent with these taxa being Pleistocene refuge species. The species are mainly monoecious, but dioecy cannot yet be ruled out in some species. In two species where sufficient material was available for investigation (*Cola chlorantha, C. roy),* the simple cymes were found to be bisexual, the terminal flower being female, the two lateral flowers often being male, a pattern previously unreported in the genus. All species are understorey shrubs or small trees of evergreen lowland or submontane forest, except *C. uloloma* of coastal semi-deciduous forest in E Africa and which is unique in showing xerophilic characteristics. The species can be divided into group A: five species mainly in E Africa with rapidly glabrescent or glabrous stems, conspicuously asymmetric leaves, few-flowered, pedunculate cymes (*C. uloloma, C. chlorantha, C. roy sp.nov., C.”udzungwa”, C. asymmetrica sp. nov.*); group B: in Lower Guinea (Atlantic coast of Africa) with hairy stems, symmetric or inconspicuously asymmetric leaves, sessile, fasciculate inflorescences: (*Cola mayimbensis, C. philipi-jonesii, C. metallica, C.moussavoui, C. stigmatosa, C. takamanda sp.nov. C. toyota sp. nov, C. “campo-ma’an”, C. zanaga sp. nov.*). Of the seven new species to science, two are named informally since the material is so incomplete (sterile), and five are formally named. Species diversity is highest in Cameroon and Tanzania, each with four species, followed by Gabon with three species. It is expected that additional new species will be found in poorly-surveyed surviving evergreen forest habitat in each of these three countries. Conservation assessments are given for each species using the IUCN 2012 standard. All species are considered threatened, with ten species being Critically Endangered (the highest category of threat), each known from a single location, with their forest habitat threatened mainly by clearance for agriculture.

## Introduction

*Cola* was included in tribe Sterculieae of Sterculiaceae *sensu lato* of the core Malvales for most of the twentieth century. Phylogenetic investigation of Malvales showed that in place of the traditional four families recognised (Malvaceae, Bombacaceae, Sterculiaceae, Tiliaceae) there is a choice of either recognising nine subfamilies in a super-Malvaceae (Bayer *et al*. 1999, Bayer & Kubitzki 2003) or recognising the same units as the families, Bombacaceae, Brownlowiaceae, Byttneriaceae, Dombeyaceae, Durionaceae, Helicteraceae, Malvaceae sensu stricto, Sparrmanniaceae (or Grewiaceae), Sterculiaceae, and Tiliaceae (Baum *et al*. 1998, Cheek & Dorr 2007, Cheek in Heywood *et al*. 2007). *Cola* can therefore now be placed either in Malvaceae-Sterculioideae or Sterculiaceae s.s. The latter approach is preferred since it is less cumbersome and minimises taxonomic instability (Cheek & Dorr 2007).

The Sterculiaceae *sensu stricto* are characterised within Malvales by unisexual flowers with a single perianth whorl which lack an epicalyx. The male flowers have an androgynophore or androphore bearing the anthers in a terminal capitulum or ring, the gynoecium vestigial and inconspicuous. Female flowers usually have a sessile or subsessile gynoecium developing into an apocarpous fruit of (1–)4–5(–15) fruitlets or mericarps, the base surrounded by indehiscent anthers. The family is pantropical, with c. 415 species arranged in 13 genera (Cheek in Heywood *et al*. 2007)

*Pterygota* Schott & Endl. pantropical, with dehiscent, woody mericarps containing dry, winged seeds, is in a sister relationship with *Cola,* while *Octolobus* Welw., confined to tropical Africa, with numerous spirally inserted indehiscent mericarps, is sister to *Pterygota*-*Cola* combined (Wilkie *et al*. 2006). The remaining genera of the *Cola* clade, *Hildegardia* Schott & Endl., *Firmiana* Marsili*, Ptercocymbium* R.Br. and *Scaphium* Schott & Endl. all have winged fruitlets and are wind-dispersed, and all but *Hildegardia* are confined to SE Asia and adjoining areas. The pantropical genus *Sterculia* L., sometimes confused with *Cola,* is in an entirely different clade, and always has dehiscent fruit with the seeds with radicle directed away from the hilum and hard-coated, borne on a placenta with irritant hairs.

The genus *Cola* with 110 – 135 species of trees and shrubs, the most species-rich genus in the Sterculiaceae, is characterised by indehiscent (rarely at length dehiscent) mericarps containing seeds with a soft, fleshy seedcoat, the radicle directed towards the hilum. The endocarp is glabrous. *Cola* is mostly confined to evergreen lowland and submontane forest in continental subsaharan Africa, with only two species in deciduous forest or woodland. While some of the species are widespread, many are extremely local, and some are known from few or single forest patches and so are vulnerable to extinction. For example, in Cameroon alone, eight species of *Cola* have been assessed as threatened (Onana & Cheek 2011). *Cola nitida* (Vent.) Schott. & Endl. and *Cola acuminata* (Pal.) Schott. & Endl. are planted throughout the tropics for their seeds which act as stimulants when chewed and are an ingredient of the eponymous and ubiquitous ‘Cola’ soft drinks. Two other species also have stimulant seeds, but are only locally cultivated (Cheek 2002a; Cheek & Dorr 2007).

Most species of *Cola* occur in Tropical Africa, with only three species, *Cola natalensis* Oliv., *Cola greenwayi* Brenan and *Cola dorrii* Cheek in South Africa (Cheek *et al*. 2018a). In East Africa (Uganda, Kenya and Tanzania), 21 species are native (Cheek & Dorr 2007). However, West and Central Africa are the heartland of *Cola*. The largest number reported for any flora region is that in the Flora of West Tropical Africa (FWTA), with 42 species, and with an additional nine imperfectly-known species (Keay & Brenan 1958). New species from West Africa have since been published (Jongkind 2004; 2013). Thirty-three species are recorded from Gabon (Hallé, 1961) and 32 from Congo-Kinshasa (Germain, 1963). The Flore du Cameroun account awaits completion. New Cameroonian species have recently been published by Kenfack *et al*. (2018), Cheek *et al*. (2020a) and for Gabon by Hallé (1988) and Breteler (2014a; 2014b). A new species has also been published from Mozambique (Cheek *et al*. 2019a). Further new species are likely to be found, particularly in Gabon, where, while 336 *Cola* specimens are recorded as being identified to species, a further 140 remain unidentified (Sosef *et al*. 2006: 395–397).

The genus was last monographed by Schumann (1900) when 33 species were recognized. Although Brenan did much research on the genus throughout its range, he confined himself, largely, to publishing accounts of new species (e.g. Keay & Brenan 1958). This paper continues preparation for a monograph of the genus that began 20 years ago (Cheek 2002a; 2002b; Cheek & Dorr 2007; Cheek *et al*. 2018a; Cheek *et al*. 2019a; 2019b; 2020a).

The majority of simple-leaved Cola species, which make up most of the species in the genus, have a distinct architecture that renders plants identifiable to genus even when sterile. During a season’s growth, pulvinate, stipulate leaves, both long and short-petioled, are produced. Once the dormant apical bud, comprised of several close-set scales, opens and the stem, first clothed with caducous leaf-scales, extends, the first true leaves produced by the now active stem apex have the largest blades and the longest petioles which may be 15 – 30 cm long. As the season advances, the leaves, both blades and petioles progressively reduce in size so that when the stem apex forms the next dormant bud for the dry season, it is subtended usually by subsessile leaves with blades that may be only 10% the area of the first-formed leaves of the season.

However, in several simple-leaved species this stage-dependent heteromorphy appears to be suppressed, the leaves during a season’s growth being all more or less the same size, and on petioles which are short, only 1 – 2 cm long. These species are less readily identifiable as *Cola* when sterile. Yet closer inspection shows that the first formed leaves of a season do have petioles longer than those produced towards the end of the growth season, although the difference is in only a few millimetres. These species retain pulvinate leaves and spiral phyllotaxy, and in other respects resemble those species described in the paragraph above.

Superficially similar to these species, also with monomorphic leaf-blades and short petioles, are several other species differing in having non-pulvinate leaves, distichous phyllotaxy, stem indumentum mainly of simple, not stellate, hairs, and male flowers with a short androphore (< 1(– 2) mm long), anthers 4 – 5 (usually not 8 – 10+), and fruits with mericarps small and globose 1 – 1.5 cm diam. (not ellipsoid to cylindrical, 4 – 30 cm long). Since this group of species appears coherent, and supported by architecture, vegetative, indumentum, flower and fruit characters, and has shown to be monophyletic in a phylogenomic analysis (unpublished), it is here formally characterised and named as a new subgenus, subg. *Disticha,* and its 14 species enumerated, including five new species which are formally described and named and a further two species which are informally named.

A separate paper reviews and revises the infrageneric classification of *Cola* (Cheek, in prep.).

## Material and methods

The patrol method was generally used to collect the specimens which form the basis of this paper (e.g. Cheek & Cable 1997). Herbarium material was examined with a Leica Wild M8 dissecting binocular microscope fitted with an eyepiece graticule measuring in units of 0.025 mm at maximum magnification. The drawing was made with the same equipment with a Leica 308700 camera lucida attachment. The number of internodes per season’s growth were measured using the position of the annual bud scale scars as reference points. Material of all species of *Cola* available at BM, EA, FHO, IEC, K, P and YA was viewed, and specimens loaned to K for study. Specimens were also inspected at the following herbaria: BR, SRGH. Flowers were rehydrated by moistening with water so that the sexual parts could be viewed. Nomenclatural changes were made according to the Code (Turland *et al*. 2018). Names of species and authors follow IPNI (continuously updated). The format of the description follows those in other papers describing new species in the genus *Cola*, e.g. Cheek *et al*. (2019a), and technical terms follow Beentje & Cheek (2003) All specimens cited have been seen unless indicated n.v. Points were georeferenced using locality information from herbarium specimens. The conservation assessments follow the IUCN (2012) standard. GeoCAT was used to calculate Red List metrics (Bachman *et al*. 2011). IUCN & UNEP-WCMC (2017) was used to check for inclusion within protected area boundaries. Herbarium codes follow Index Herbariorum (Thiers, continuously updated).

## Taxonomic Results

*Cola subgenus Disticha* Cheek subg. nov.

Type species: *Cola philipi-jonesii* Brenan & Keay (1955)

Monoecious, possibly sometimes dioecious, small evergreen shrubs, less usually small trees. Phyllotaxy distichous; petioles short, < 10 mm long, pulvini not formed; stage dependent leaf heteromorphy absent; dormant buds with bud scales present. Inflorescences bracteate, 1-flowered, fasciculate or simple, few-flowered cymes, sometimes pedunculate. Flowers dull white, yellow-white or green, perianth divided by 70 – 90% of its radius into 5 – 6 lobes, lobes with white membranous margin or not, outer surface usually with stellate hairs, inner surface finely papillate, otherwise glabrous (rarely with some stellate hairs. Male flowers with androphore <1(– 2) mm long, anthers 4 – 6 in a single whorl. Female flower with carpels 4 – 5, styles erect, linear (except *C. stigmatosa,* flattened), at the apex spreading. Fruits with mericarps globose < c. 1 cm diam., indehiscent, 1-seeded (where known), red or orange, smooth, lacking surface sculpture. Seedcoat thick, white, fleshy.

### DISTRIBUTION & HABITAT

14 species in SE Nigeria, Cameroon, Gabon, Republic of Congo, DR Congo, Kenya, Tanzania, Malawi. Lowland or submontane evergreen, rarely semi-deciduous forest on limestone (*Cola uloloma*). Species are concentrated in:

A. the Eastern Arc Mountains and Coastal Forests (EACF) of Tanzania and Kenya which contains three species: *Cola uloloma, C. roy, C. “udzungwa”*.
B. the Cross-Sanaga Interval of SE Nigeria and western Cameroon with four species (*Cola philip-jonesii, C. takamanda, C. metallica, C. toyota*).

These two biogeographic areas are well-known for their high levels of endemic plant species, many of which are considered relicts and thought to be connected to refuge areas (Skarbek 2008; Gereau *et al*. 2016; Cheek *et al*. 2001). The Cross-Sanaga includes the areas with the highest species and generic diversity per degree square in tropical Africa (Barthlott *et al*. 1996; Dagallier *et al*. 2020), rivalled only by the EACF of Tanzania. Outside of these two focal areas, *Cola chlorantha* occurs in W. Tanzania and northern Malawi, and, outside of the Cross-Sanaga, Cameroon also holds *Cola “campo-ma’an”* sited within one of the putative Pleistocene Refugia (Sosef 1994), as are the remaining species of the Lower Guinea phytogeographic area, respectively: *Cola mayimbensis* (Belinga Mts), *C. stigmatosa* (Nyanga), *C, moussavoui* (Ngounie) all of Gabon, and *C. zanaga* (Chaillu Mts, Republic of Congo) and *C. asymmetrica* (Mayombe Mts, D.R. Congo).

It is notable that not a single record of a single species occurs in the Congo Basin, and that the subgenus is absent from West Africa (west of the Cross River). This is consistent with the subgenus being a refuge taxon since the Congo basin lost its forest in the Pleistocene arid periods, except for the swamp areas of the river Congo and its major tributaries. No species of the subgenus is recorded from swamp forest.

Other groups of species occurring in EACF and Lower Guinea, including the Cross-Sanaga include the genera *Zenkerella* Taub. and *Kupea* Cheek & S.A. Williams (Leguminosae and Triuridaceae respectively; Cheek *et al*. 2003; Cheek 2004)

While most species (nine) can be considered lowland (below 800 m. alt) and no species occurs in montane forest (above 2000 m alt.) five species occur in submontane forest (c. 800 – 2000 m alt.). At 1750 – 1830 m, *Cola chlorantha* attains the highest altitude, followed by *C. roy* (1500 m). *Cola mayimbensis*, *C. toyota* and *C. “udzungwa”* also reach or surpass 1000 m elevation.

### CONSERVATION STATUS

All seven of the species known before this paper was written have been assessed for their IUCN extinction risk status, and all seven have been found to be highly threatened (Endangered, EN or Critically Endangered, CR) apart from *C. uloloma* (Vulnerable, VU), and it can be argued that this last species should be re-assessed as EN (see under that species). The five new species to science formally published in this paper are all considered Critically Endangered because they are single site endemics that face threats, except for *C. zanaga,* with two locations, considered EN. Of the two Imperfectly Known species, *C. “campo-ma’an”* would likely rate CR, while *C. “udzungwa”,* because it is in a National Park currently considered well protected, would likely be assessed when published formally as VU. Until species have been formally published, it is difficult to have formal assessments accepted by IUCN (Cheek at al. 2020b). In summary, more than half the species (eight) accepted in this paper are considered Critically Endangered, four Endangered and two Vulnerable. The priority species for attention must be the three species known from single gatherings in unprotected areas and nor recorded in more than 60 years. These must be at the highest risk of global extinction. Global extinctions in tropical Africa are increasing. Apart from those recorded in Humphreys *et al*. (2019), additional global plant species extinctions have been recently recorded in the areas where the taxa treated in this paper are concentrated: the Cross-Sanaga Interval (Cheek & Williams 1999; Cheek *et al*. 2018b; 2018c; 2019c), in the EACF of Tanzania (Cheek & Luke 2022) and in Gabon (Moxon-Holt & Cheek 2021; Cheek *et al*. 2021a). This makes it imperative to seek measures to protect these highly threatened *Cola* species e.g. by inclusion in Tropical Important Plant Areas programmes (Darbyshire *et al*. 2017) to reduce the risk of their global extinction.

### VERNACULAR NAMES AND USES

Very few of the species have local names and uses, consistent with the fact that they are so rare that they have not come to the attention of local communities. The exception is the most widespread and common species, *Cola uloloma*, which has been used to make rough beds (see that species). However, it is likely that the fruit of all species is edible, with a thick brittle, sweet seedcoat as in the majority of *Cola* species. Yet, because the fruits of our taxon are so small compared with those of the rest of the genus, they are unlikely to be specially sought after by humans.

### NOTES

Two species groupings can be recognised in the subgenus:

Group A comprising *Cola uloloma, C. roy, C. chlorantha, C. asymmetrica* and presumably *C. “udzungwa”.* These all have glabrous stems (rarely with a few hairs near the apical bud), pedunculate inflorescences, and a tendency to asymmetric leaf-blades. Apart from *Cola asymmetrica*, they are confined to E Africa.

Group B, comprising *Cola philip-jonesii, C. takamanda, C. toyota, C. metallica, C. mayimbensis, c. moussavoui, C. stigmatosa C. zanaga,* (and presumably *C. “campo-ma’an”*) have hairy stems, glabrous stems, sessile inflorescences or single flowers, and leaf asymmetry is absent or inconspicuous. These species all occur in the Lower Guinea Phytogeographic domain of the western coast of Africa.

However, these groupings are based on observations of limited material, and may have little value. They should be tested by a molecular phylogenetic study.

Of the 14 species recognised in this paper, complete material, with both complete male and female flowers and fruits, was available for only two species: *Cola philipi-jonesii* and *C. uloloma.* All the remaining species lack one or more of these important structures, and in the case of the two Imperfectly Known species (see end of species sequence), all three structures. Complete flowers remain unknown for three other species (*Cola asymmetrica, C. metallica* and *C. toyota*).

There are no recorded observations of pollinators in any of the species, but day-flying insects are suspected to be the likely pollinators. The flowers of *Cola uloloma* have been reported as “sweet-smelling”. Seed dispersal has also not been recorded, but primate dispersal is well known in the genus, and although small, the fruits of our taxon have the features of the majority of the genus. However, the high proportion of range-restricted species suggests that seed dispersal is not effective at extending range-size in most species.

Many *Cola* species are considered dioecious, and in some cases this is heavily supported by evidence (e.g. Cheek *et al*. 2018a). However, investigation of the (usually limited) flowering material for each species (where available) in this paper suggests that in this taxon the species are mainly, sometimes, or always monoecious (see the individual species accounts). Investigation of two of the species which have pedunculate 3-flowered cymes (group B) showed that the terminal flower is often female, while the two later-produced lateral flowers of the cyme are males (see below under *C. chlorantha* and *C. roy*), a pattern previously unrecorded in the genus. However, in a third species of group B, *Cola uloloma,* this pattern was not detected. Instead, the majority of flowers were found to be male with a ratio of 22 male: 1 female flower, but the sample was limited by the material available.

Species of *Cola* subg. *Disticha* when first viewed, even by experienced botanists, are often misplaced in families outside Malvales, unless the interior of the flowers (if present) is examined. This is because they lack the standard vegetative identification characters of *Cola* (see introduction), and even the indumentum can be dominated by simple hairs, not the stellate hairs that help diagnose Malvales. Species included in this genus have been identified as *Cleistanthus* (Euphorbiaceae), as “?*Dovyalis”* (Flacourtiaceae, now Achariaceae), as Menispermaceae and as *Strombosia* (Olacaceae). It is perfectly possible that additional specimens, even additional species unknown to science, await discovery in herbaria where they are misidentified under these or other families.

#### Identification key to the species of *Cola* subgenus *Disticha*

Note that the two incompletely known *Cola* species A (*Cola “*campo-ma’an”) and B (*Cola “*udzungwa”) are not included in the key since fertile material is lacking.

1. Leaf-blades conspicuously asymmetric, wider by several mm on one side of the midrib than the other; inflorescences shortly pedunculate, cymose; stem glabrous (rarely, *C. roy*, with a few stellate hairs near the apical bud). 2

1. Leaf-blades ±symmetrical, if asymmetric, inconspicuously so; inflorescences sessile and fasciculate or 1-flowered, rarely seemingly cymose (*C. toyota*) and then sessile; stem with distinct vestiture, 10 – 100% covered in simple and/or stellate hairs persistent for at least the first internode and usually several internodes more 5

2. Flowering stems becoming white-waxy several internodes below apex; flowers white, yellow or pinkish brown, with tepal lobes spreading (perpendicular to the main axis), margins hyaline, white, undulate. Coastal Kenya & Tanzania **1. *C. uloloma***

3. Flowering stems green or purple, matt, not developing a waxy surface; flowers green (where known) with tepal lobes ascending, tepal margins as wide as, and concolorous with the rest of lobe 3

4. Leaves with (3 –)4 – 5(– 7) secondary nerves on each side of the midrib; (4.4 –)5 – 8 x (1.5 –)1.9 – 2.8(– 3.2) cm; 5 – 10-inflorescences per stem. W. Tanzania…**2. *C. roy sp. nov*.**

2. Leaves with 5 – 9 secondary nerves on each side of the midrib, 10 – 13.8(– 15) x (2.2 –)3.9 – 5(– 5.4) cm; 1 – 2-inflorescences per stem 4

3. Leaves with 5 – 7 secondary nerves on each side of the midrib; petioles c. 20% covered in simple hairs; pedicel length 5 – 7 mm. Western D.R.C **3 *C asymmetrica sp. nov*.**

4. Leaves with 7 – 9 secondary nerves on each side of the midrib; petioles glabrous; pedicel length 2 – 3 mm. Malawi & Tanzania… **4. *C. chlorantha***

5. Leaf acumen spatulate - the distal ⅕ wider than the proximal part; stem densely covered in white stellate hairs which slough off as scurf, revealing a grey-white epidermis, lenticels absent or inconspicuous. S.E.Nigeria ***C. philipi-jonesii***

6. Leaf acumen narrowly triangular or ligulate, widest at base; stem indumentum solely or mainly of simple hairs, sometimes with a small fraction of stellate hairs; epidermis glossy purple, dark brown or black, with white lenticels conspicuous. Cameroon, Gabon, Congo. 6

7. Leaf blade with (8 –)10 – 13(– 15) lateral nerves on each side of the midrib; inner surface of perianth lobes with stellate hairs; margin of lobes undifferentiated, not white or crinkled or hyaline; stigmas large, flat, pendulous, reaching anthers in female flowers.Gabon… **6. *C. stigmatosa***

5. Leaf-blade with 6 – 8(– 9) lateral nerves on each side of the midrib; inner surface of perianth lobes lacking hairs; margin of perianth lobes white, crinkled, hyaline; stigmas erect & filiform, if flat then short, not pendulous 7

6. Leaf-blades drying glossy metallic dark grey or metallic blackish green below (abaxial surface). S. W. Region, Cameroon **7. *C. metallica***

7. Leaf-blades drying green, rarely black, matt, not metallic below (abaxial surface). 8

8. Stipules persistent, densely simple hairy; flowers pendulous; perianth campanulate (perianth lobes erect); androphore 2 mm long. Gabon **8. *C. mayimbensis***

9. Stipules caducous, or, if moderately persistent, glabrous; flowers erect; perianth with lobes spreading; androphore 0.1 – 1.3 mm long… 9

10. Leaf-blades 11.7 – 20 x 3 – 7 cm, galls absent 10

8. Leaf-blades 8 – 11 x 2.5 – 3.9 cm, galls present 11

9. Leaf-blades oblanceolate (rarely obovate-elliptic); pedicel 10 mm long. Republic of Congo **9. *C. zanaga sp. nov*.**

10. Leaf-blades elliptic; pedicel 3 – 3.5 mm long… ***C. moussaoui***

11, Leaf-blades with (7 –)8 – 9 lateral nerves on each side of the midrib; stipules extremely caducous (unknown); stem 10 – 20% covered in hairs; 1 inflorescence per stem; pedicels 2.5 – 4.5 mm long. C. Region, Cameroon **11. *C. toyota sp. nov*.**

11. Leaf-blades with 6(– 7) lateral nerves on each side of the mid-rib; stipules moderately persistent; stem 30 – 40% covered in hairs; numerous inflorescence per stem; pedicels 5 – 7 mm long. S W Region, Cameroon **12. *C. takamanda sp. nov*.**

#### 1. Cola uloloma

Brenan (1956:150; Beentje 1994:163; Cheek & Dorr 2007:36). Type: Tanzania “Tanganyika Territory: Pangani District, Bushiri Estate, fl. no date, fl. *Mrs Helen Faulkner* 654 (Lectotype selected here, see note below, K barcode K000241032!; isolectotypes BR barcodes BR0000006290382 BR000000629034!, K barcode K000241033!)

**DISTRIBUTION & HABITAT.**S.E. Kenya and N.E. Tanzania, coastal semi-deciduous forest sometimes on limestone, with *Julbernardia magnistipulata* commonly dominant*, Craibia brevicaudata* a common understory tree, and *Mkilua fragrans*, *Craterogyne kameruniana* and *Uvariodendron kirkii* common in the shrub layer, also in the canopy *Paramacrolobium, Cynometra, Inhambanella, Manilkara, Erythrophleum, Hymenia, Malacantha Chlorophora*, *Odyendea, Bombax, Trachylobium, Scorodophloeus, Combretum schumannii, Carpodiptera, Cassipourea, Cordyla, Pleurostylia, Haplocoelopsis, Cola pseudoclavata, C. minor, Ricinodendron, Drypetes.* Lianes include *Combretum, Salacia, Dichapetalum*; 30 – 500 m alt.

**ADDITIONAL SPECIMENS. KENYA. K7, Kwale District:** Mwele Mdogo Forest, Shimba Hills, 12 miles S.W. of Kwale, fr. 23 Aug 1953, *Drummond & Hemsley* 3972 (EA, K!); Kwale District, Muhaka Forest, st. 2 March 1977, *Faden* 77/608 (EA, K!); Mangea Hill Top, st. 8 Apr 1987, *Luke, W.R.Q. & Robertson* 307 (EA, K!); Kaya Kambe, fr. 9 July 1987, *Robertson & Luke, W.R.Q.* 4787 (EA, K!); Pangani Rocks, near stream through limestone rocks, fls 10 July 1987, *Luke, W.R.Q. & Robertson* 494 (EA, K!); Kaya Ribe, on sandstone, around Mleji and Mbunzi Rivers, buds. 10 July 1987, *Robertson & Luke, W.R.Q.* 4843 (EA, K!); Buda Mafisini Forest Rsv, st 23 Feb 1989, *Luke, W.R.Q. & Robertson* 1681 (EA, K!); - Marenji Forest Rsv., st. 26 Feb 1989, *Luke, W.R.Q. & Robertson* 1756A (EA, K!); Shimba Hills, Mwele, fr. 16 Oct. 1991, *Luke, W.R.Q.* 2934 (EA, K!); Kaya Muhaka, fl. 9 June 1994, *Luke, W.R.Q. & Gray* 3983(EA, K!); Dzombo Hill, st. 7 Feb 1989, *Robertson, Beentje Luke, Q., Khayota* 206, (EA, K!); **Kilifi District**: Kaya Kivara, st. 7/8 July 1987, *Robertson & Luke, W.R.Q.* 4760 (EA, K!). **TANZANIA.** Pangani District, Bushiri Estate, fl. no date, *Mrs Helen Faulkner* 654 (EA, K); Tanga Prov., Pangani, Mwera, Mwanamgaru, Kwa Besa, fl. 28 March 1957, *Tanner* 3462 (DSM, EA, K!)

**CONSERVATION STATUS.** Beentje (1994:163) lists *Cola uloloma* as “Vulnerable, ?Endangered” and as collected at Pangani, Buda, Muhaka, Marenji, Gongoni and Mangea. In Cheek & Dorr (2007: 37) it is stated “this species seems rare, poorly known and restricted to a few small coastal forest fragments, mostly the Kaya forests of Kenya. Its extent of occurrence is estimated as less than 20,000 km^2^. Given that forest quality is declining at several of these patches, and that less than ten locations based on 14 specimens are known, *Cola uloloma* is here assessed as VU B2a,b(iii), i.e. vulnerable to extinction*”.* This statement overlooks the fact that because so much habitat between the surviving forest patches has been lost, this species is “severely fragmented” within the definition of IUCN (2012). If the area of occupancy is under 500 km^2^ as expected, the species should be assessed as EN B2ab(iii). No formal IUCN assessment has been published of this East African species although it is hoped that an account will be published in future. (Luke pers. comm. to Cheek). Since 2007 no additional sites are known to have been discovered for this species. Numerous other threatened species occur within the range of *Cola uloloma* including, in Kenya alone, *Ancistrocladus robertsoniorum* J. Léonard (Ancistrocladaceae, Léonard 1984; Cheek 2000), *Afrothismia baerae* (Thismiaceae, Cheek 2004), *Lukea quentinii* Cheek & Gosline (Annonaceae, Cheek *et al*. 2022a) and *Vepris robertsoniae* Q.Luke & Cheek (Rutaceae, Cheek & Luke 2022)

**PHENOLOGY.** Flowering March, June, July; fruiting July-Sept.

**ETYMOLOGY.** Unknown.

**LOCAL NAMES & USES.** Mnofi (Zigua) *Tanner* 3462 (K!) “used for making rough beds”

**NOTES.** *Cola uloloma* is sympatric at several sites with *C. minor* Brenan and *C. pseudoclavata* Cheek (formerly misidentified as *C. clavata* Mast., Cheek & Dorr 2007)

Two sheets (sheet 1 and sheet 2) are labelled as holotype at K. Sheet 1 is selected here as lectotype because it has the original field notes cited in the protologue and because it is of better quality than sheet 2. The collector recorded that the plants were dioecious: only male flowers were seen but both fruit and male flowers are present on the same stem (monoecious) in *Luke & Gray* 3983(K). However, male flowers appear to heavily outnumber females. Searching open flowers on this specimen for female flowers, only male flowers were encountered until the 23^rd^ flower proved to be female. Faulkner noted that the flower colour was “pinky brown” which is not usual in the subgenus, where cream, off-white or green is more normal. Yet they are described also as “cream” (*Luke & Gray* 3983) and “white/yellow with brown hairs” *(Luke & Robertson* 494, K). According to *Tanner* 3462, the flowers are “yellow-brown, aromatic”.

Brenan (1956) in the protologue, points directly to the affinities of his new species not being with any other in E. Africa, but with *C. mayimbensis* of Gabon and with a Nigerian specimen then in press as *Cola philipi-jonesii.* In fact, *Cola uloloma* was the third species to be published, and the first from E. Africa, of subgenus *Disticha* characterised in this paper.

Mrs. Helen Faulkner (1888 – 1979) collector of the type material, made over 5,000 herbarium collections in Tanzania, continuing into her ninetieth year. Most of her collections were made in the triangle between Tanga, Handeni and Pangani and included valuable material from the rapidly diminishing natural habitat on limestone and from forest relics (Polhill & Polhill, 2015).

*Cola uloloma* was initially known from a single collection, *Faulkner* 654 collected at unknown date in N.E. Tanzania (Brenan, 1956), it was not until 1953 that it was discovered in S.E. Kenya, when fruit were found (*Drummond & Hemsley* 3972). Today it has been recorded in more sites in Kenya than in Tanzania. *Cola uloloma* is the only species of subg. *Disticha* recorded from semi-deciduous forest (all other species are from evergreen forest) and it seems to have thicker leaves, perhaps an adaptation to this drier habitat. It is unique in the subgenus in that the flowering stems, initially glabrous and green, develop a white, waxy covering at the fourth or fifth internode below the stem apex, and ovoid dormant axillary buds are present on most specimens, also putative xerophilic traits. The petioles are glabrous apart from a few stellate and simple hairs on the adaxial surface.

A specimen from the Pugu Hills, Tanzania, identified as this species, *Hawthorne* 1645 (FHO, K!) would extend the range south of that which is currently accepted, but in the absence of flowers that identification is treated with caution since it lacks the white waxy epidermis that helps characterises the taxon.

Sterile material can resemble and has been mistaken for *Cleistanthus* (Euphorbiaceae). Sterile material collected from the Udzungwa Mts of Tanzania as this species are identified here as *Cola ‘udzungwa’* see Imperfectly Known Species at the end of this sequence. The Udzungwa specimens lack the white waxy epidermis and have scattered red sessile glands on the abaxial blade surfaces, not seen in *C. uloloma*.

Perhaps because of its apparent adaptation to a drier habitat, unique within the subgenus, *Cola uloloma* although rare, threatened and restricted to about eight forest patches, is by far the most widespread and common of all species of the subgenus.

#### 2. Cola roy

Cheek sp. nov. Type: Tanzania, Iringa, Mufindi, Lulanda, 8°36’S, 35°37’E, 1500m, afromontane rainforest, fl. 24 Nov. 1988, *Gereau & Lovett* 2559 (holotype K! barcode K000511990, isotypes DAR, MO). (Fig. 1)

**Fig. 1.**
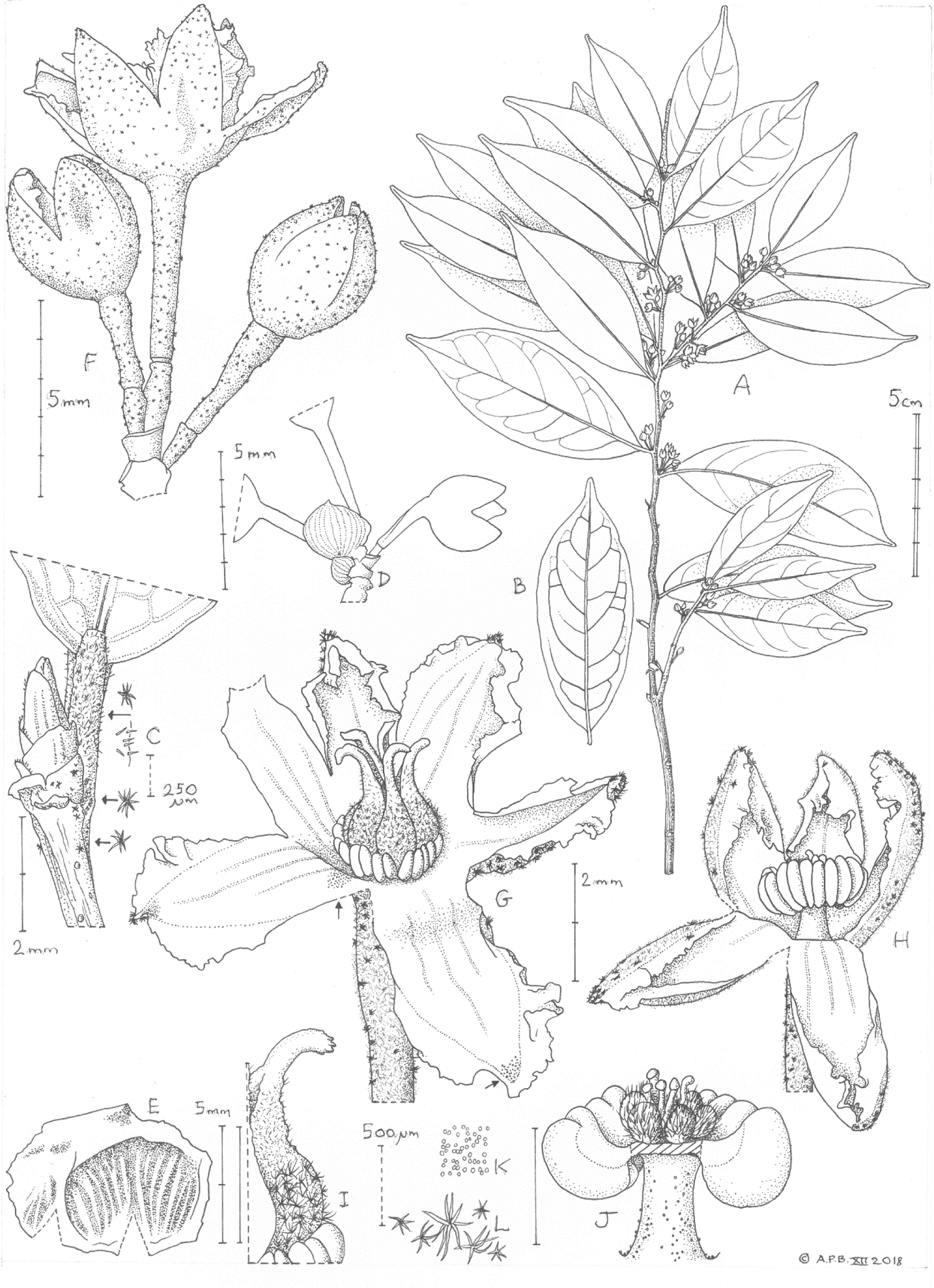
*Cola roy* **A** habit, flowering branch; **B** abaxial surface, large leaf; **C** apical bud at stem apex with leaf base, inset showing details of hairs from (top to bottom) petiole, bud-scale, and stem; **D** inflorescence sketch showing caducous bracts still in position; **E** inflorescence bract, inner surface; 3-flowered inflorescence with terminal female, and two lateral male flowers; **F** female flower, perianth tube cut and one lobe pulled down to show pistil; **H** male flower, tepal tube cut and two lobes pulled down to display androecium; **I** detail of ovary showing a carpel with style and stigma, from G**; J** detail of male flower from **H** with anther thecae removed to show gynobasic pistillodes; **K** detail of papillae from adaxial surface of perianth lobes; **L** trichomes from abaxial surface of the female perianth lobes, larger hairs are dark red, smaller ones are colourless. All drawn from *Gereau & Lovett* 2559 (K) by ANDREW BROWN

*Monoecious*, *evergreen tree* 5m tall, trunk 10 cm diam. at 1.5m above ground. Features of the trunk not recorded. Leafy stems terete, 1 – 2 mm diam. at lowest leafy node, internodes (0.5 –)0.8 – 1.2 cm long (Fig. 1A) epidermis brown, glossy, smooth in current season, ridged in previous season, lenticels dull white, raised, circular 0.3 – 0.8 mm diam. on previous season’s growth, inconspicuous, glabrous apart from a few stellate hairs on the distal. internode, hairs 0.15 – 0.2 mm diam., 7 – 9-armed, highly caducous and inconspicuous. *Terminal buds* narrowly ovoid, 2.1 – 4 x 0.9 mm, glossy, olive brown, bud-scales 6 – 8, (Fig. 1C) outermost transversely ovate-oblong, wider than long, c. 0.8 x 1.8 mm, innermost lanceolate, c. 1.8 x 0.7 mm, apex rounded, margin fringed with minute simple hairs, dorsal surface with 2 – 3 shallow longitudinal ridges. *Leaves* (3 –)5 – 8 over the length of a season’s growth, apparently persisting for two seasons, last produced leaf of a season about 5/8ths the length of that first produced; blades discolorous, upper surface drying glossy dark brown, lower surface pale matt brown-green, narrowly elliptic (end of season’s leaves) to oblong-elliptic (first produced blades), (4.4 –)5 – 8 x (1.5 –)1.9 – 2.8(– 3.2) cm, acumen short, broad, apex rounded, rarely tapering and acute, 0.25 – 0.5(– 0.75) cm long, sides shallowly convex at the broadly acute base. Secondary nerves (3 –)4 – 5(– 7) on each side of the midrib, yellow, arising at 70 – 80° from the midrib, arching steadily upwards towards the margin, in the distal two-thirds of the blade the nerves often connecting with the secondary nerve above via a short tertiary nerve, forming a looping nerve c. 2 mm from the margin, domatia absent; tertiary and quaternary nerves forming a reticulum, glabrescent, with sparse, inconspicuous red, stellate hairs, 8 – 12-armed, 0.15 – 0.3 mm diam. on the secondary nerves. *Petioles* terete 2 – 5 mm long, 0.7 mm diam., drying highly crinkled, adaxial surface 50 – 80% covered in simple, erect, colourless hairs 0.05 – 0.08 mm long, (Fig. 1C), stellate hairs very sparse, 7 – 9-armed 0.15 – 0.2 mm diam., abaxial surface glabrous; pulvini absent. *Stipules* caducous, not seen. *Inflorescences* shortly pedunculate, cymose, 3(– 4)-flowered, 1 per axil, at 5 – 10 successive nodes, the terminal flower of each inflorescence female, opening first, the lateral flowers male, opening afterwards; buds not seen; peduncle 0.9 – 1 x 1.2 – 1.3 mm; rhachis 0.7 – 0.8 x 0.9 – 1.0 mm; bracts orbicular, concave, with 7 – 15 parallel longitudinal ridges, 1 – 2 x 1 – 3.5 mm (Fig. 1E), distal bracts longer than proximal, glabrous, caducous. *Flowers* each 5 – 9 mm diam., opening in succession within inflorescences, while stem apex is dormant, from stems of current and previous season’s growth. *Pedicels* articulated about ¼ the length from the base, 3.6 – 6.4 mm long, basal part (below articulation) 1.1 – 1.7 mm long, distal part (above articulation) 2.5 – 4.7 mm long (Fig.1D), 0.75 mm diam., narrowest (0.5 mm diam.) at base of pedicel, and widest (1 mm) at apex, indumentum of dimorphic stellate hairs a) large dark red 8 – 10-armed, 0.3 – 0.5 mm diam.; b) small, translucent, 6-armed, 0.1 – 0.2 mm diam., pedicel about 10 – 15% covered, almost entirely in type b) hairs (Fig.1L). *Perianth* green, divided by 2/5 to ½ into 5(– 6) green ascending lobes, each oblong-triangular 2.5 – 3.2 x 1.6 – 2.1 mm, apex acute with a short, densely red stellately hairy mucro, margins uneven in width, distal half with wide white, membranous, undulate-lacerate flanges, 0.7 – 0.8 mm wide, proximal half margins slender; margins inflexed in bud, spreading at anthesis, tube shallow bowl-shaped, 1.3 – 1.4 x 3 – 3.25 mm, inner surface of perianth with three parallel longitudinal lines, surface densely covered, with translucent vesicles (Fig. 1K) globular, 0.02 – 0.03 mm diam., 400 – 500 per mm ², outer surface of perianth 10 – 15% covered in hairs as in pedicel, denser, 20 – 30% cover on lobes, densest towards apex. *Male flowers* smaller than females, c. 5 mm diam. at anthesis, perianth lobes 5(– 6), androphore cylindrical 0.6 – 0.85 mm long, 0.4 – 0.5 mm wide, widening gradually to 0.7 mm wide at the base, surface glabrous, thinly scattered with minute translucent vesicles; anthers uniseriate 4(– 5), glabrous, in a disc 0.8 – 0.9 mm long, 1.9 – 2.2 mm diameter; ovary vestigial mostly concealed within the anther head (Fig. 1H), carpels 5, ellipsoid, 0.3 – 0.4 x 0.25 – 0.3 mm, 50 – 60% covered in simple, patent hairs 0.1 – 0.2 mm long; styles gynobasic, filiform, erect, 0.4 – 0.5 mm long, glabrous; stigmas capitate (Fig. 1J). *Female flowers* c. 9 mm diam., perianth lobes 5 – 6, gynophore absent; anthers 5, in ring at base of ovary, same dimensions as in male fowers; ovary ovoid 2.8x 2 mm, conspicuously divided into 5(– 6) carpels. Carpels each erect, narrowly ovoid, 1.1 – 1.2 x 0.5 – 0.7 mm, outer surface densely white stellate hairy (70 – 90% cover), hairs 0.1 – 0.2 mm diam., arms 5 – 6 erect or spreading. Styles erect c. 1 x 0.2 mm, tapering gradually from the ovary, proximal 2/3 with adaxial groove, slightly warty, c. 10% covered in patent simple hairs 0.05 mm long; distal 1/3 cylindrical, glabrous, spreading or slightly reflexed. Stigma punctiform or slightly verrucate (Fig. 1 G&I). Fruits and seeds unknown.

**RECOGNITION.** differing from *C. chlorantha* F. White in the smaller ((4.4 –)5 – 8 x (1.5 –)1.9 – 2.8(– 3.2) cm), more closely-spaced leaf internodes (0.5 –)0.8 – 1.2 cm long, with short, broad acumen 0.25 – 0.5(– 0.75) cm long, (vs leaves (8.5 –)9 – 11.8 x (3.1 –)3.5 – 4.2(– 4.3) cm, internodes (1.2 –)1.7 – 2.2 cm long, acumen 1.3 – 1.7 cm long, tapering gradually to an acute point. For additional differential characters see Table 1.

**Table 1.** Characters distinguishing *Cola roy* from *Cola uloloma* and *Cola chlorantha* (data for the last two species mainly from Cheek & Dorr (2007)).

|  | *Cola uloloma* | *Cola roy* | *Cola chlorantha* |
| --- | --- | --- | --- |
| Habitat & altitudinal range | Semi-deciduous coastal forest; 30 – 500m | Submontane forest; 1500m | Submontane forest; 1500 – 1730m |
| Geographic range (FTEA divisions in brackets) | Kenya & Tanzania (K7, T3,?6) | Tanzania (T7) | N. Malawi & Tanzania (T4) |
| Adaxial petiole densely simple patent hairy | Yes | Yes | No |
| Colour and surface of leaf-blade, adaxial surface | Pale green, matt | Dark brown, glossy | Green, matt |
| Androphore length | 1.5 – 2 mm | 0.6 – 0.85 mm | Not available |
| Perianth colour (excepting white lobe margins) | White, yellow or pinkish brown | Green | Green |
| Perianth lobe posture at anthesis | Patent or slightly reflexed | Ascending | Ascending |
| Leaf-blade dimensions | 4 – 12 x 1.7 – 4.7 cm | (4.4 – )5 – 8 x (1.5 – )1.9 – 2.8( – 3.2) cm | (8.5 – )9 – 11.8 x (3.1 – )3.5 – 4.2( – 4.3) cm |
| Lateral nerve number each side of midrib | 5 – 7 | (3 – )4 – 5( – 7) | 7 – 9 |
| Internode length | (0.4 – )0.8 – 1.8 cm | (0.5 – )0.8 – 1.2 cm | (1.2 – )1.7 – 2.2 cm |
| Acumen shape and length | Usually long, tapering to an acute apex, 0.5 – 1.8 cm long | Short, broad, apex broadly rounded, 0.25 – 0.5( – 0.75) cm long | Long, tapering gradually to an acute point, 1.3 – 1.7 cm long |
| Perianth tube shape, dimensions | Cup-shaped 1.25 – 1.5 x 2 – 2.5 mm | Shallowly bowl-shaped 1.3 – 1.4 x 3 – 3.25 mm | Bowl shaped c. 3.5 x 3 – 5 mm. |

**DISTRIBUTION & HABITAT**. Tanzania: Iringa, Mufindi Escarpment, Lulando (or Lulanda) forest; submontane forest; 1500 m alt.

**ADDITIONAL SPECIMENS.** None are known.

**CONSERVATION.** Only a single collection is known of *Cola roy*, despite searches of the East African Herbarium (EA) and that of Tanzania (DAR) for additional collections. The grid reference for the specimen locality given on the label is probably read from a map and not a Global Positioning System (GPS). This is deduced from a) the date, 1988 – when such devices were rarely available for botanical survey work, and b) since only whole degrees and minutes are given: unlikely if a GPS has been deployed. An error margin of several hundred metres can therefore be deduced. Visiting the geolocation given on Google Earth takes us to a highly anthropic landscape of patched field and plantations of tea (*Camellia sinensis* L.) and conifers (for timber). However, about 200 m SE of the geolocation is a polygon measuring 0.89 x 0.5 km that appears largely uncultivated and to possibly have been a former forest reserve which is suspected to be the “Lulando Forest” of Darbyshire (2009): “For a site of such limited extent, Lulando Forest has a rich biodiversity and represents one of the few intact submontane rainforest sites in the Southern Highlands. It is also the southernmost site for several Eastern Arc faunal endemics” (Darbyshire 2019)

The central part of this forest patch still appears to be natural, if degraded forest, extending 0.65 x 0.16 km (Google Earth imaging viewed 31 Dec 2018). This is the likely source of the collection since no other patches of natural forest occur nearby. From the irregular boundary of the surviving forest and the variation in tree canopy size, it appears that this patch is being gradually reduced in area and in quality, probably due to wood extraction. It appears not to be protected. Therefore, we assess *Cola roy* as Critically Endangered CR B1+B2ab(iii) since area of occupancy and extent of occurrence are both 4 km² using the IUCN preferred cell-size of that area, since there is a single location with clear threats. We hope that by drawing attention to this species by delimiting and naming it, efforts might now be made to protect it at its only known location near Lulando, Mufindi and to find additional locations where it might survive. Additional range-restricted species unique to this location are *Isoglossa ventricosa* I. Darbysh. (Darbyshire 2009) and *Coffea lulandoensis* Bridson (1994). Bridson stated: “Recent exploration of the Lulando Forest has revealed many new taxa,. and shown this forest to be especially rich in endemics”.

**PHENOLOGY.** Flowering in November.

**ETYMOLOGY.** The specific epithet commemorates Roy Gereau, staff taxonomic botanist of Missouri Botanical Garden (MO), U.S.A. the lead collector of the only known specimen, who has had a long interest in the taxonomy and conservation of tropical African plants and their conservation, particularly in Tanzania where he has lead MO’s programmes for many years. He is also commemorated by the genus *Gereaua* Buerki & Callm. (Sapindaceae, Buerki *et al*. 2010).

**NOTES.** The type, and only known specimen of *Cola roy*, *Gereau & Lovett* 2559, bears the field determination “Flacourtiaceae, *Dovyalis*?” indicating how difficult it can be to recognise these short-petioled species as belonging to the genus *Cola*. The specimen was determined by Roy Gereau in 1998 as *Cola uloloma* Brenan, to which it is similar and probably related. I redetermined the specimen as *Cola chlorantha* F. White in 2002, and the specimen is cited under that name (and not *Cola uloloma)* by me in Cheek & Dorr (2007), on the basis that *Gereau & Lovett* 2559 also has green flowers (not whitish to pinkish brown as in *C. uloloma*) derives from 1500m altitude (not 30 – 500m), has inconspicuous bracts (not conspicuous) and ascending perianth lobes (not patent). However, I overlooked that it is morphologically discordant in several characters from *Cola chlorantha*, especially in the much smaller, more closely spaced leaves which dry glossy dark brown on the upper surface and have only a short, broad, rounded acumen. These vegetative features alone immediately separate it from the three known specimens of *Cola chlorantha*. Other features that separate *Cola roy* from *C. uloloma* and *C. chlorantha* are shown in Table 1.

In Cheek & Dorr (2007:37) I noted several specimens that had been brought to my attention by Quentin Luke as being referable to *Cola uloloma* or to *Cola chlorantha*, all of which were at EA and which I had not seen. These included *Luke* & *Luke* 8747, *Luke et al.* 11311, both also from T7. These specimens, and others previously unstudied, were obtained on loan from EA to determine whether or not they might represent additional specimens of this sp. nov. All were ruled out, on vegetative morphology alone. Instead, the specimens mentioned are referred to a separate taxon: see ¬Imperfectly known species B’, *Cola ‘udzungwa’,* at the end of the taxonomic sequence.

#### The bisexual cymes of *Cola roy*

While one small group of *Cola* species has racemose inflorescences, and another paniculate inflorescences (represented by *Cola nitida* and *Cola gigantea* A. Chev., respectively) the remainder, the majority of species, have fasciculate inflorescences, or single-flowered inflorescences.

However, close inspection of the inflorescences of *Cola roy* reveals that although they appear fasciculate, a peduncle is present (Fig. 1F). Remarkably the terminal, first developed flower, is female, while the two (rarely three) later produced flowers of the cyme, are all male. This pattern appears not to have been documented previously in *Cola*, where most species are dioecious, and where monoecy has been recorded, no such pattern has been noted. A similar pattern has been found in *Cola chlorantha* (see under that species), but not the geographically close and probably closely related *C. uloloma* although that species is also monoecious.

#### 3. Cola asymmetrica

Cheek sp. nov. Type: “Congo Belgica (now Democratic Republic of Congo), Mvula, plateau de la Ngundu et de la Kingulu”, alt. ±500m, fr. 20 Jan. 1949, *Toussaint* 753 (holotype K barcode K000874908; isotypes BR n.v.). (Fig. 2).

**Fig. 2.**
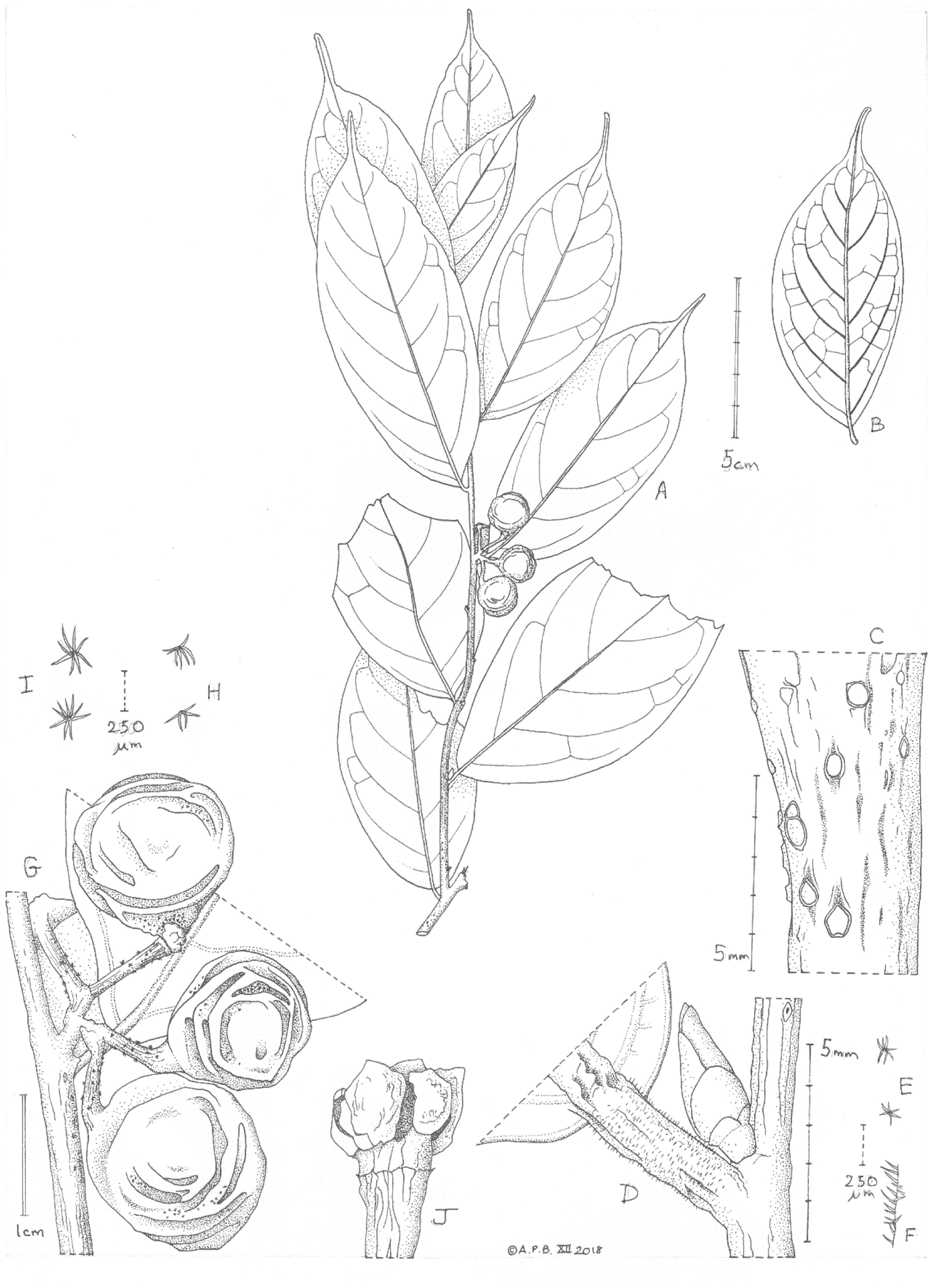
*Cola asymmetrica.* **A.** habit, fruiting, leafy stem; **B.** leaf, abaxial surface; **C.** older stem detail showing lenticels; **D.** junction of leaf base with stem showing axillary bud; **E.** stellate hairs from axillary bud-scales; **F.** detail of simple hairs on petiole; **G.** detail of infructescence from A; **H.** stellate hairs from pedicel shown in G; **I.** stellate hairs from fruit surface in G; **J.** head of fruiting pedicel bearing receptacle with mericarps removed, showing stamen remains (densely dotted) between the mericarp cicatrices. All drawn from *Toussaint* 753(K) by ANDREW BROWN.

*Evergreen shrub* (probably) height and trunk unknown. *Leafy stems* terete, 3-5 mm diam. at lowest leafy node, green to brown-purple, finely longitudinally ridged, lenticels raised, bright white, orbicular or longitudinally slightly elliptic, (0.4 –)0.6 – 0.7 mm diam., glabrous; internodes 1.5 – 3.3 cm long; scale-leaves not seen. *Terminal and axillary buds* pale brown, narrowly ovoid c. 4.2 x 1.5 mm, bud-scales c. 6, closely appressed, glossy, glabrous excepting a few 6 – 7-armed stellate hairs 0.15 – 0.2 mm diam. at base of outermost scales (Fig. 2E). *Leaves* 4 – 9 over the length of a season’s growth, persisting for two seasons or more, drying pale grey-green, leathery, asymmetrically elliptic, 10 – 13.8(– 15.2) x (2.2 –)3.9 – 4.8(– 5.4) cm, the side of the blade (measured from midrib to margin) proximal to the stem apex 1 – 4 mm wider than that on the side distal to the stem apex, (Fig. 2B), acumen (0.9 –)1 – 1.8 x 0.2 – 0.4 cm, base asymmetrically obtuse or rounded, sides convex, midrib raised, yellow, keel-like on adaxial surface; secondary nerves 5 – 7 on each side of the midrib, yellow, arising at c. 50° from the midrib, arching steadily upwards towards the apex, united in the distal half to third by a looping nerve 3 – 5 mm from the margin, domatia absent; tertiary and quaternary nerves forming a raised reticulum visible on the abaxial and adaxial surfaces, glabrous. *Petioles* terete, 4 – 6 mm long, 1 – 1.5(– 1.8) mm, wrinkled and yellow when dry, c. 20% covered in simple, erect, colourless hairs 0.08 – 0.15 mm long (Fig. 2F); pulvini absent. Stipules caducous, not seen. *Inflorescences* known only at fruiting stage, 4 – 5-flowered, 1 per axil, 1 per leafy stem at third to sixth node from the stem apex; peduncle 2 – 3 mm long, partial-peduncles c. 2 mm long; pedicels 5 – 7 mm long, 0.8 – 1 mm diam., articulation absent; 5 – 10% covered in stellate hairs 0.1 – 0.15 mm diam., arms erect, 5 – 7 (Fig. 2H). *Flowers* known only at fruiting stage with perianth fallen, stamens 5, alternating with carpels (Fig. 2J). Carpels 5. *Fruit* carpels 1 – 5, each 1-seeded, globose 1 – 1.5 cm diam., apex rounded, style remains absent or inconspicuous, base sessile, rounded; surface drying orange, lacking grooves or sculpture, indumentum of 8 – 9-armed, 0.3 mm diam., stellate hairs covering 40 – 50% of the surface, caducous.

**RECOGNITION.** Differing from all other *Cola* Sect. *Disticha* species in Central Africa in the glabrous stems (not mainly simple hairy), leaf-blades strongly asymmetric, one side much wider than the other (not symmetric or very slightly asymmetric), a single fertile node per leafy stem, bearing 4 – 5 fruits (all other species have multiple fertile nodes except *C. metallica* Cheek, and that has but 1(– 2) fruits per node).

**DISTRIBUTION & HABITAT.** Democratic Republic of the Congo, the Mayumbe Mts (deduced, see Notes below). Lowland evergreen forest (deduced); c. 500m alt.

**ADDITIONAL SPECIMENS.** None are known.

**CONSERVATION.** The single deduced (see Notes below) location for *Cola asymmetrica* is outside the Luki Biosphere Reserve, in a formally unprotected area dominated by evergreen forest. The area of occupation is estimated as 4 km^2^ using the preferred IUCN grid-cell size, and extent of occurrence as the same. However, using the historic imagery function of Google Earth, it is clear that between the most recent available imagery there (7 April 2015, Landsat/Copernicus, accessed 18 June 2022) and imagery there from March 2003, about 50% of the forest present in 2003 had been lost by 2015 in an approximately 4 km x 4 km square including the deduced specimen point. This rapid loss, apparently through clearance for agriculture, over just 12 years indicates that the species, if it still exists, will become extinct at its sole known location within a few years. *Cola asymmetrica* is here assessed as Critically Endangered (CR B1ab(iii)+B2ab(iii) based on the data above. It is to be hoped that the species will be found at additional locations in the Mayumbe mountains, e.g. the large and adjacent Luki Biosphere Reserve. Sadly however, Google Earth imagery suggests that forest habitat inside Luki, affected by dense settlements through much of its area, may be no more protected than that surviving forest outside the Reserve.

**ETYMOLOGY.** Named for the strongly asymmetric leaf blades.

**NOTES.** *Toussaint* 753 was distributed from BR as *Strombosia* (Olacaceae). The Kew sheet is annotated in pencil as “?*Cola*” in the hand of Frank White. Examination of the c. 250 unidentified *Cola* specimens at BR failed to uncover additional specimens of this taxon. The species is not included in the Flore de Congo Belge account for *Cola* (Germain 1963). There is no indication of habit or habitat on the sheet at K. Searches of the specimen database of BR for this specimen failed to locate the expected duplicate there. It is possible that only a single sheet exists and that was distributed to K in error. The specimen is not present on gbif.org at the time of writing (July 2022).

The specimen label is likely to be erroneous as to the location data: “Mvula, plateau de la Ngundu et de la Kingulu”. Kingulu is located at 5° 39’ 29.77”S, 18° 12’ 43.31”E, at 763 m alt. in gallery forest in predominantly grassland (viewed on Google Earth). However, the specimen tag (put on when collected as is normal practice) number *Toussaint* 753 is not consistent with this location. This is for two reasons: 1) specimen numbers of *Toussaint* (from gbif.org) below and above 753 are from evergreen forest at Gimbi, Mayumbe at 5° 31’S, 13° 22’E, e.g. *Toussaint* 750 collected 19 Jan. 1949 (*Justicia biokoensis* V.A.W. Graham (Acanthaceae) and 763 collected 21 Jan. 1949 at Gimbi, vallée de la Mvusi (*Adenia bequaertii* subsp. *occidentalis* W.J. de Wilde, Passifloraceae); 2) the collection date given on the label of the type specimen, 20 Jan. 1949 is consistent with the number series, being the day after 750 was collected, and the day before 763 was collected. It seems unlikely that *Toussaint* having collected several days or weeks in the Mayumbe Mts, would travel 536 km nearly due East, across the cataracts of the mighty Congo river, to a remote site, collect a specimen there, and return the same day, the same distance to Gimbi, Mayumbe to continue collecting there for several days more. Logistically this would have been nearly impossible then, as it is now, and even if it were possible, it is unlikely. The specimen concerned may have been misplaced. Therefore, it can be concluded beyond reasonable doubt that the collection location was Gimbi, vallée de la Mvusi, in the Mayumbe Mts rather than that given on the specimen label.

The Mayumbe Mts (also known as the Maiombe, or Mayombe), clothed in evergreen forest (where not cleared for agriculture), of altitude rarely exceeding 600m m elevation, extend 400 – 500 km, more or less parallel to the Atlantic coast c. 50 – 150 km inland from immediately N of the Congo River in DRC, northwards through Cabinda (the exclave of Angola), the Republic of Congo, into Gabon. They mark the division between the coastal plain and the higher inland plateaux. Numerous other rare and threatened plant species are restricted to the Mayumbe Forests e.g. *Nepthytis mayombensis* de Namur & Bogner (Araceae, Bogner & de Namur 1994).

#### 4. Cola chlorantha

**B.** F. White (Dowsett-Lemaire & White, 1990:83; White, Dowsett-Lemaire, Chapman 2001:556; Cheek & Dorr 2007:37). Type: Malawi, N. Region, North Viphya, Choma Mt Forest, near Muzuzu, fl. 7 Nov. 1982, *Dowsett-Lemaire* 471 (holotype K barcode K000241027!; isotype BR barcode BR628932!).

**DISTRIBUTION & HABITAT.** Malawi and Tanzania; evergreen east African rift submontane forest; 1500 – 1750 – 1830 m alt. Dowsett-Lemaire in Hyde *et al*. (2020), describes the forest at the Musisi Hill location as with canopy 25 – 30 m in height, with large gaps. *Chrysophyllum gorungosanum* is the commonest large tree, with *Macaranga kilimandscharica* (above 1700 m), and *Parinari excelsa* and *Polyscias fulva* also common.

The midstratum is dominated by *Garcinia kingaensis* and *Brucea antidysenterica* the understorey. *Cola chlorantha* is in the understorey at 1750 – 1830 m, with *Podocarpus falcatus* and *Croton megalocarpos.* At the type location, Choma Forest, *Chrysophyllum* is again the commonest larger tree (canopy is 25 m in height) followed by *Grewia (Microcos*) *mildbraedii.* Here *Cola chlorantha* was widespread in the understorey to a height of 10 – 15m (Dowsett-Lemaire in Hyde *et al*. (2020)).

**ADDITIONAL SPECIMENS. TANZANIA,** Kigoma Dist., Kasakati, fl. Sept. 1965, *Suzuki* B-45(K!); **MALAWI** “SF1033: Musisi Hill”, *Dowsett-Lemaire* ecol. 116 (FHO).

**CONSERVATION STATUS.** *Cola chlorantha* was assessed as Endangered (EN B1ab(iii)) by Lawrence & Cheek (2018) using data on the Malawi populations from Dowsett-Lemaire (pers. comm. 2018). At the type location, Choma Mt, where it was reported “not uncommon” in the protologue, the main forest patch, only 800 m long and 70-80 m wide, has become considerably thinned and is vulnerable to fires invading from adjoining land (Dowsett-Lemaire in Hyde *et al*. 2020). At Musisi Hill the main forest consisted of two patches of 73 and 114 ha. In Malawi most forest lacks formal protected area status, and even forest reserves have been cleared of forest: the species may not survive in Malawi. In Tanzania, at the Mahale Mts, in the last 5 – 10 years the forests are now reported to be protected, and fire-breaks are installed. However, the species has not been recorded there since 1965 (Lawrence & Cheek 2018). Since 1965 much forest at the locality has been lost and it is not certain that the species survives in Tanzania. Precise co-ordinates are unavailable for all records since they pre-date availability of Global Positioning Systems. *Cola chlorantha* is believed not to be in cultivation nor in a seedbank. It is advisable that attempts are made to refind this species in the field and to ensure its protection through all possible measures.

**PHENOLOGY.** Flowering in September (Tanzania) and November (Malawi).

**ETYMOLOGY.** From the Greek, referring to the green flowers.

**LOCAL NAMES & USES.** None are known.

**NOTES.** The type specimen at K had been distributed from FHO as *Cola greenwayi* Brenan, but Brenan (16 July 1987) annotated it as “Cola? sp. nov. Certainly not *C. greenwayi* because of the short petioles uniform in length. Its affinity is with *C. uloloma* Brenan (E. Africa) but venation and leaf-base different, and flowers apparently larger. Please get lots more material of both sexes!” The only isotype, at BR (not at FHO as indicated in the protologue) is stamped with an FHO registration number but has been marked “ex”.

I included in *Cola chlorantha Gereau & Lovett* 2559 (K!) from Lulanda Forest, Mufindi, Tanzania (Cheek & Dorr 2007:37). This was an error since that specimen is now the basis of *Cola roy* Cheek (see above). The three East African species of subgenus *Disticha, C. uloloma, C.chlorantha* and *C. roy* are discussed under the last species. The differences between these species are given in the key and in Table 1.

*Cola chlorantha* is very incompletely known. Fruit are lacking. More work is needed to compare the two, far-disjunct (800 km separation) populations in Tanzania and Malawi. The Tanzanian material has a purple stem epidermis with white lenticels while that of Malawi is green with flaking bark. It may be that they are separable taxonomically.

Françoise Dowsett-Lemaire, a keen field biologist, is the only person known to have seen this species alive in Malawi. Apart from the type location at Choma Mt, she also observed it at Musisi Hill (White *et al*. 2001: 556) and collected saplings (leaves narrower and more attenuate than of mature leaves) as an ‘ecological specimen’. Although this was stated in the protologue to be deposited at FHO it is thought to have been transferred to BR as was the isotype.

*Cola chlorantha* is monoecious. Examination of 16 flowers at 7 nodes on the holotype revealed that, while the 3-flowered cymes at two nodes were all female, at another the terminal flower was female and the two laterals male. At three of the remaining nodes, only two flowers were observable of the three in each cyme. In two of these one of the pair was a male and the other female, while in the third pair both were females. The single flower at the seventh node was female.

#### 5. Cola philipi-jonesii

Brenan & Keay (Keay 1954:329; Brenan & Keay 1955: tab.3534.) Type: Nigeria, “Ogoja Province, Ikom Division, Afi River Forest Reserve, about 0.8 km west of River Nwop along south boundary of Boje enclave, in high forest, 16 May 1946, *A.P.D. Jones & Onochie* FHI 5838 (Lectotype K, barcode K000240899; isolectotypes: K barcode K000240898, BR barcode BR0000006290399; FHI n.v.; P barcode P00368371).

**DISTRIBUTION & HABITAT.** Only known from the type location in S.E. Nigeria close to the border with Cameroon. Lowland evergreen forest.

**ADDITIONAL SPECIMENS.** Brenan & Keay (1955) state that “a specimen numbered 72388 in the Imperial Forestry Institute herbarium (FHO) at Oxford bears leafy shoots which may belong to the new species. The date, place and collector are unknown. It is stated that “The label and the detached fruits on the sheet do not belong.” It has not been possible for the author to view this specimen.

**CONSERVATION STATUS.** *Cola philipi-jonesii* is assessed as Endangered EN B1+2c (World Conservation Monitoring Centre 1998). “Unprotected forest has been almost completely felled up to the paths boundaries” it is stated. Other threats given are cultivation of annual and perennial non-timber crops, livestock farming and ranching, and logging and wood harvesting. However, at the type locality in 1946, the collectors recorded that it was a “very common understorey shrub.” Yet it has not been recorded by scientists since that time. Targeted searches at the type locality are needed to confirm that the species survives and to fill the many gaps in knowledge that exist for this species.

According to Imong & Wood (2007) the Afi River Forest Reserve covers approximately 380 km² at the headwaters of the Afi River in the northern part of Cross River State. It is an area of lowland forest and ridge forest that is rarely used by Cross River gorillas but is seen as an important corridor connecting other areas with larger populations: the Mbe Mountains and the Afi Mountain Wildlife Sanctuary. Imong & Wood (2007) reported the Forest Reserve to be neglected, resulting in an increase in logging, farming and hunting. Approximately three farms and three signs of logging were recorded per km of transect within the reserve, and they stated that they could see the entire reserve becoming farmland in the foreseeable future unless action is taken (Imong & Wood 2007). This does not bode well for the global survival of *Cola philipi-jonesii*.

Given the single location, an AOO and EOO estimated as 4 km^2^, and the threats indicated, the extinction risk of *Cola philipi-jonesii* is assessed here as Critically Endangered (CR B1ab(i-iii)+B2(ab(i-iii).

**PHENOLOGY.** Flowering and fruiting in May.

**ETYMOLOGY.** Commemorating a Mr. A.P.D. Jones “one of the collectors of this and many other plants in Nigeria, in recognition of the considerable amount of work that he did towards a revision of *Cola*. His untimely death in 1946 deprived West Africa of an outstanding botanist and us of a personal friend.” (Keay 1954: 329)

**LOCAL NAMES & USES. N**one known.

**NOTES.** *Cola philipi-jonesii* is the most western and only known Nigerian species of subgenus *Disticha*. Since it was the first described species in the areas with the distinctive morphology of subg. *Disticha*, all specimens of the adjoining Cameroonian species of the subgenus were initially compared with it and often first identified as it. From all of these *Cola philipi-jonesii* differs in the spatulate acumen (the apex distinctly wider than the proximal part) and the stems which initially are densely stellate (not simple) hairy and which when the hairs are shed are pale grey-white, matt, not dark glossy purple.

There are two sheets labelled ‘Holotype’ at K. The second, labelled ‘sheet 2’ is here selected as second stage lectotype (barcode K000240899) since it has flowers, and the other sheet appears sterile.

*Cola philipi-jonesii* is suspected to be monoecious because both female and male flowers are known from the single gathering. In addition, the authors of the species did not indicate dioecy in the protologue.

The only known location of *Cola philipi-jonesii* at Afi River is host to several other rare, range-restricted or point-endemic species, such as *Placodiscus glandulosus* Radlk. (*Jones & Onochie* in FHI 18633, Sapindaceae, Keay 1958:329*), Saxicolella flabellata* (G. Taylor) Cheek (based on *Keay* FHI 28240, Podostemaceae, Cheek *et al*. 2022b), and *Ledermanniella tenuifolia* (G. Taylor) C. Cusset, (*Keay* FHI 28241, Podostemaceae, Cusset 1987). The adjacent Oban Hills also contains several point endemics such as *Anchomanes nigritianus* Rendle (Araceae, Moxon-Holt & Cheek 2021), not seen for over 100 years.

The discovery of *Cola philipi-jonesii* was part of a major botanical survey of the forests of Nigeria (and of the then British Cameroons) in the 1950’s apparently initiated and masterminded by RWJ Keay (Cheek *et al*. 2000: 27)|, of the Forest Research Institute, Nigeria at Ibadan which houses the major herbarium FHI (Thiers, continuously updated). Keay published eight other new *Cola* species from Nigeria in partnership with Brenan (IPNI, continuously updated).

#### 6. Cola stigmatosa

Breteler (2014: 105). Type: Gabon, Nyanga, c. 41 km on Tchibanga-Mayomba Rd., 3° 4.6’S, 10° 44.1’E, 21 Oct. 2009, *Bissiengou, Breteler, Niangadouma & Boussiengoi* 418 (holotype WAG barcode WAG0324295!; isotypes BR, G, K, LBV, MA, MO, P, WAG barcode WAG0324294!).

**DISTRIBUTION & HABITAT.** Gabon, only known from the type collection; lowland evergreen forest; c.100 m alt.

**ADDITIONAL SPECIMENS.** None are known.

**CONSERVATION STATUS.** The habitat of *Cola stigmatosa* at its single known location is severely threatened by ongoing extensive slash and burn agriculture, consequently the species was assessed as Critically Endangered (CR B1ab(i,ii,iii) + 2ab(i,ii,iii)) by Baldwin & Cheek (2019).

**PHENOLOGY.** Flowering in October; fruiting unknown.

**ETYMOLOGY.** Referring to the relatively large, conspicuous stigmas (Breteler 2014:105).

**LOCAL NAMES & USES. N**one known.

**NOTES.** *Cola stigmatosa* is remarkably distinct from all other known species of subgenus *Disticha.* Although it has leaf-blades similar in shape and dimensions to other species, the number of secondary nerves is much higher, (8 –)10 – 13(– 15) on each side of the midrib, rather than the 5 – 7 which is usual. The flowers are also remarkable because the stigmas are large, flat and descend around the ovary partly concealing it and reaching the anthers. In all other species of the subgenus the stigmas are erect, or slightly curved, much smaller than the ovary, never concealing it, and never descending. While the tepal margins of most other species have a white, undulating membranous band, this is absent in *C. stigmatosa* the tepal margin being undifferentiated in colour and texture. The inner surface of the tepals of *C. stigmatosa* are stellate hairy – in all other species where flowers are known, the inner surface lacks hairs.

*Cola moussavoui* is suspected to be monoecious because both female and male flowers are known from the single gathering. In addition, the authors of the species did not indicate dioecy.

As with *Cola moussavoui* this name appears not yet to be registered on the Naturalis website that holds WAG specimens, and it was not possible to view images (accessed 19 June 2022). However, the protologue illustration is excellent. Equally it was not possible to access records of the isotypes at other herbaria probably because duplicates are still in the process of being distributed.

Gabon has 32 species of *Cola* recorded, almost equalling Cameroon in diversity (Sosef *et al.,* 2006: 395). However as pointed out by Breteler (2014) many more are likely to be discovered. One hundred and 38 specimens of the genus remain undetermined to species from Gabon (Sosef *et al*. 2006).

#### 7. Cola metallica

Cheek (Cheek 2002: 409; Cheek *et al.,* 2004: 186; Onana & Cheek 2011: 332 – 333).Type: Cameroon, S.W. Region, Nguti, near the Banyang Mbo Wildlife Sanctuary, Research station path to sanctuary via Nlowoa and Mbu river crossings, fr. 25 Nov. 2000, *Cheek* 10598 (holotype K000691491!; iso YA!). *Cola* sp. nov. 1 aff. *philipi-jonesii* Brenan & Keay, *sensu* Cheek in Cable & Cheek (1998:134).

**DISTRIBUTION & HABITAT.** Cameroon, only known from S.W. Region around Mt Cameroon, northwards to Banyang Mbo Reserve at Nguti. Lowland evergreen forest; c. 50 – 700 m alt.

**ADDITIONAL SPECIMENS. CAMEROON.** Mt. Cameroon, Batoke, Mile 9, fr. 4 June 1992, *Kwangue* 97 (K!, SCA); between Upper Boando and Etome, fr. 9 Sept 1992, *Kwangue* 132 (K!, YA) ibid 8 Dec. 1993, *Faucher* 31 (K!, SCA); Upper Boando fr. 30 Nov. 1993, *Cable* 266 (SCA!); ibid 1 Dec. 1993, *Cable* 284 (K! SCA); Bomana, Onge Forest, c. Oct. 1993, *Ekema* 591 (K, SCA); Ekombe-Mofato fr. 23 May 1994, *Watts* 1169 (K!, SCA, YA); Kupe-Muanenguba Division, Bakossi Forest Reserve, fr. 23 Oct. 1998*, Etuge* 4299 (K!, YA); Nguti, near the Banyang Mbo Wildlife Sanctuary, Research Station path to sanctuary via Nlowoa and Mbu river crossings, fr. 25 Nov. 2000, *Cheek* 10598 (holo K!; iso YA!).

**CONSERVATION STATUS.** *Cola metallica* was assessed as CR A1c+2c in Cable & Cheek (1998:134), updated to CR A2c (Cheek *et al*. 2004:186; Onana & Cheek 2011:332), and to CR A4c (Cheek & Lawrence 2018). The species is known from four locations, at each of which there are threats to the habitat due to clearance of forest for agriculture, plantations and logging. Due to the scale of habitat loss, more than 80% of the population has and is being lost over a three generation period from 1990 to 2050 it is estimated. Since the species was published in 2002, no additional records of it have been made.

**PHENOLOGY.** Fruiting May, June, Sept.-Dec.; flowering season unknown.

**ETYMOLOGY.** Referring to the metallic appearance of the leaves of herbarium specimens which dry glossy metallic dark grey or metallic blackish green on the lower (abaxial) surface (Cheek 2002: 409).

**LOCAL NAMES & USES.** None is known.

**NOTES.** *Cola metallica* is highly conspicuous when in fruit because these are bright orange, contrasting with the glossy green leaves. Despite this fact and that it has been collected in fruit 6 months of the year it is extremely rare, being infrequent within its small range within S.W. Region. It cannot be mistaken for any other species in the section because of the metallic dark grey or black-green colour of the dried abaxial leaf-blade. All other species dry green, or yellow green. Female flowers, and intact male flowers, are unknown. The species is monoecious because fruiting specimen were found to have the remains of old male flowers attached to them.

*Cola metallica* was one of three new species to science of *Cola* that came to light (Cheek, 2002) as a result of botanical surveys on and around Mt. Cameroon (Cheek 1992, Thomas & Cheek 1992, Cheek *et al*. 1996) which culminated in The Plants of Cameroon, a conservation checklist (Cable & Cheek 1998). Additional rare and threatened species discovered to be new to science in these surveys include *Octoknema mokoko* Gosline & Malécot (Gosline & Malécot 2012), *Oxygyne duncanii* Cheek (Cheek *et al*. 2018c), *Impatiens etindensis* Cheek & Eb. Fischer (Cheek & Fischer 1999), *Impatiens frithii* Cheek (Cheek & Csiba 2002), *Ancistrocladus grandiflorus* Cheek (Cheek 2000) and most recently *Drypetes burnleyae* Cheek (Cheek *et al*. 2021b). Mount Cameroon is a major centre of diversity for *Cola,* with 24 species (Cheek 2002a), but also for other plant groups e.g. achlorophyllous mycotrophs (Cheek & Williams, 1999).

Several of the first specimens collected were not identified in the field as *Cola*, but as Menispermaceae, presumably because they so little resemble *Cola* vegetatively and because of the apocarpous fruits.

#### 8. Cola mayimbensis

Pellegr. (Pellegrin 1950: 189; Hallé 1961:66).Type: Gabon, Lastoursville, Mayimba, fl. 12 Dec 1929, *Le Testu* 7765 (Lectotype selected here P00368369!; isolectotypes P00368368! P00368367! BM000645978!; BR628950!; BR628953!; HBG51228!; K000540739 photo!; M02201433!; M03208598!, WAG0003373!).

**DISTRIBUTION & HABITAT.**Gabon, known from locations at Belinga and Lastoursville. Lowland and lower submontane forest 400 – 1000 m alt.

**ADDITIONAL SPECIMENS.GABON**, *Lastoursville,* fl. 15 Dec. 1929, Le Testu 7643 (P06655622! P06655623!); Belinga, fl. 16 Nov. 1964, N. Hallé 3183 (P00730168!); ibid. fl. 21 Nov. 1964, *N. Hallé* 3289 (P06655629!; Belinga, Mines de fer, fr. 19 July 1966, *N. Hallé & Le Thomas* 93 (P06655630!); ibid Relevé 1.66 Belinga st., no date, *N. Hallé & Le Thomas* 594 (P06655627!) ibid, Belinga, Relevé 1 A8 666. st. July 1966. *N. Hallé & Le Thomas* 666 (P06655624!).

**CONSERVATION STATUS.** *Cola mayimbensis* was assessed as Endangered (EN B1ab(iii)+2ab(iii)) by Cheek & Baldwin (2019a). It is threatened at both its two locations. At Belinga by open-cast iron ore extraction which for the moment is on hold, and at Mayimba, near Lastoursville, by slash and burn agriculture (Cheek & Baldwin 2019a).

**PHENOLOGY.** Flowering in November and December; fruiting in July.

**ETYMOLOGY.** Meaning ‘from Mayimba’ the name of the village and area whence the type specimen was collected in Gabon.

**LOCAL NAMES & USES. N**one are known.

**NOTES.** Georges Le Testu (1877 – 1967) collector of the type material of *Cola mayimbensis* was a French colonial administrator who was also a prolific collector of plant specimens, primarily in Gabon. Most of the endemic Gabonese *Cola* species were based on his specimens and published by Pellegrin (1950) including *Cola letestui* Pellegr., *Cola tsandensis* Pellegr., *Cola mahoundensis* Pellegr., *Cola lissachensis* Pellgr., and *Cola glaucoviridis* Pellegr.

**A.** *N. Hallé & Le Thomas* 666(p) is sterile, and is perhaps a juvenile plant since it bears leaves much larger than in the flowering specimens. It also bears a second label, *N. Hallé* 32898, probably due to a mix up. Nicholas Hallé who first collected *Cola mayimbensis* at Belinga, had previously authored the Flore Du Gabon account of Cola (Hallé 1961).

*Cola mayimbensis* (not to be confused with *Cola mayumbensis* Exell), was the first species to be published of subgenus *Disticha*, predating the second, *C. philipi-jonesii*, by five years.

Until surveys were conducted at Belinga for the planned and controversial future open-cast iron ore mine in the mid 1960’s, the species was only known from the type location at Lastoursville. At Belinga it was recorded in submontane (cloud) forest at altitudes of 800 m, 900 m, 950 m and 1000 m (*N. Hallé & Le Thomas* 93, P), while at the type locality c. 400 m elevation is recorded. *Cola mayimbensis* and *Cola metallica* are the only species of the subgenus *Disticha* in Lower Guinea which are known from more than one location and which have multiple rather than single, collections.

*Cola mayimbensis* is remarkable among the W-C African species of the subgenus for three features – firstly the tepal lobes do not spread (as in all other species of subg. *Disticha*), but remain erect, giving the perianth a campanulate shape, and secondly the flower is pendant in life (*Hallé 3289,* P) not erect as in other species. Thirdly, the androphore is 2 mm long, twice the length of other species (where the androphore length is usually 1 mm or less). These two features may be linked, and may support a pollination syndrome different from those of other species of the subgenus for which flowers are known.

It is not clear whether the species is monecious or dioecious. The most detailed investigation of the species by *Hallé* (1961) is not conclusive on the matter. Examination of all seven flowers on one specimen showed all to be male suggesting dioecy, but male flowers can outnumber females in monoecious species (see *Cola uloloma*), and a single specimen with so few flowers is insufficient to conclude the matter.

References to *Cola mayimbensis* outside of Gabon are believed not to be based on correctly identified specimens, but rather to other species of subgenus *Disticha* enumerated in this paper.

Lectotypification. Three sheets of the type number are present at P. Here the lectotype is selected (P00368369) because it alone has the location and date of collection data that are cited in the protologue, suggesting that it was directly used by Pellegrin. It is also of as good quality and representative of the species as the other duplicates.

#### ’9. Cola zanaga

Cheek sp. nov. Type: Republic of Congo (Congo-Brazzaville), Lekoumou Prefecture, between Moutienne and Mt Lebayi, midridge on motorable track, site of plot 3 of March 2009, fl. 14 October 2009, *Cheek* 15853 (holotype K000616409; isotypes IEC! WAG!). (Fig. 3)

**Fig. 3.**
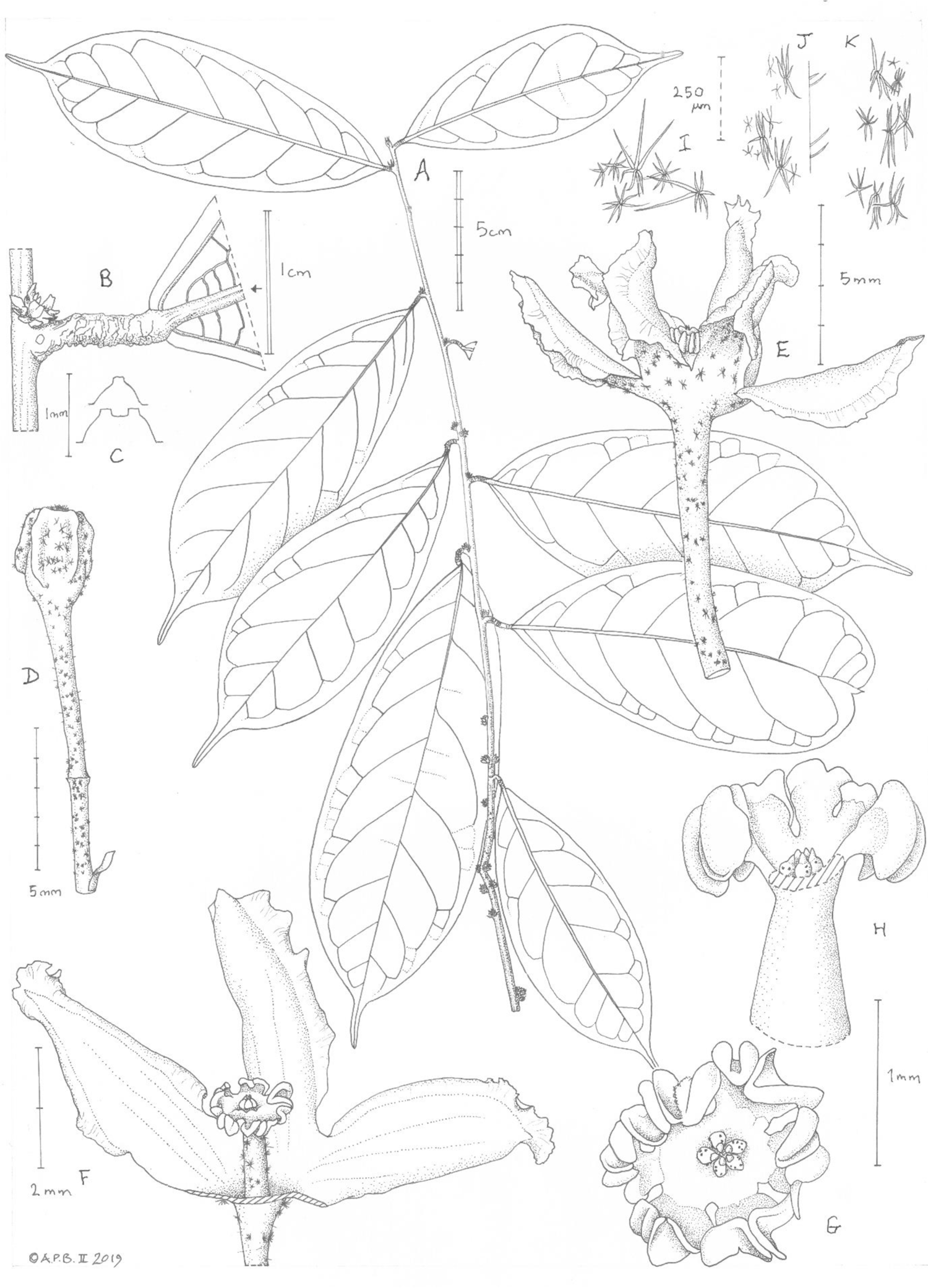
*Cola zanaga* **A** habit, leafy stem with inflorescences (flowers fallen); **B** node with axile fasciculate inflorescence (flowers fallen), and petiole and base of the leaf-blade**; C** transverse section if midrib abaxial surface of leaf-blade; **D** flower bud; **E** male flower, side view; **F** male flower, two perianth lobes removed to show androphore and anther-head; **G** anther-head, plan view; **H** atypical glabrous androphore, anther-head with two stamens removed to show vestigial carpels; **I** indumentum from outer surface of perianth; **J** indumentum complement from base of pedicel, with simple hairs shown in profile; **K** indumentum immediately below pedicel articulation. **A-K** from *Cheek* 15853 (K). Drawn by ANDREW BROWN.

*Dioecious, evergreen shrub or tree* 2m tall. Leafy stems terete, 2 – 2.5 mm diam. at lowest leafy node, internodes (0.8 –)1.9 – 3.0(– 3.5) cm long (Fig. 3A) phyllotaxy distichous, alternate, epidermis dull dark purple-brown, longitudinally ridged, lenticels on nodes distant from the apex, sparse, bright white, orbicular 0.5 mm diam.; 10 – 20% covered in erect, simple, colourless hairs 0.025 – 0.1 mm long; scale leaves absent or inconspicuous. *Terminal buds* ovoid, 4 – 5 x 2 – 2.5 mm, bud-scales 5 – 6, naviculate-elliptic, 3 – 3.5 x 1 – 1.2 mm, at first densely stellate hairy, soon glabrescent, glossy, brown, indurate. *Leaves* 9 – 10 per 14 – 25 cm of stem (the presumed length of a season’s growth), persistent, drying mid-green on both surfaces, blades oblanceolate, obovate-elliptic (10 –)11.7 – 17.8 x (4 –)4.7 – 6(– 6.2) cm, acumen ligulate to narrowly triangular, apex rounded, (0.7 –)1.5(– 1.8) x 0.3 cm, secondary nerves (5 –)7(– 8) on each side of the midrib, bright white (herbarium specimens), arising at c. 50° from the midrib, arching up, becoming parallel to the margin and uniting directly or via a subsidiary nerve to the secondary nerve above, forming a looping nerve c. 5 mm from the margin, domatia absent, tertiary and quaternary nerves forming a fine inconspicuous reticulum, glabrescent. *Petioles* terete (2.5 –)5 – 10 mm long, drying yellow-green, highly crinkled, nearly glabrous except for a few stellate hairs on abaxial surface, simple hairs on adaxial surface as stem (Fig. 3B), midrib raised, with a single or double flat-topped ridge (Fig. 3C). *Stipules* highly caducous, present only at stem apex, subulate, 4 x 0.8 mm, glabrescent hairs simple, present only where unabraded, margin scarious. *Inflorescences* axillary, fasciculate, initiated at second node below apex, increasing in size with age, 1 – 2-flowered near stem apex, increasing to 7 – 8-flowered on burrs at 18 – 19 nodes below apex, burrs c. 4 x 6 mm. Bracts increasing in size and degree of bifurcation towards apex, basal bract ovate 0.8 x 0.8 mm, acuminate, second bract ovate 1.2 x 0.9 mm, apex minutely bifurcate, third bract obovate 2 x 1 mm, bifurcation 0.25 mm; fourth bract lanceolate, 4 x 1.5 mm, bifurcate for 2 mm, soon glabrescent, glossy brown, indurated. Pedicel 10 mm long, articulated 4 mm from the base (Fig. 3D), 15 – 20% covered in a mixture of 1) dimorphic copper stellate hairs, a) larger hairs, arms directed along axis of pedicel, 0.35 – 0.4 mm long, 6-armed; b) smaller hairs (base of pedicel only) arms radiating symmetrically, 0.06 – 0.12 mm diam., 4 – 6-armed, and 2) purple-black stellate hairs, hairs (5 –)6 – 8-armed, smaller hairs 0.15 – 0.2 mm diam., arms equal in length, larger hairs 0.5 – 0.6 mm diam., arms unequal, those at one side 2 – 4 x as long as the others. *Perianth* (5 –)6-lobed (Fig. 3E), tube broad and shallow, c. 1 x 4 mm; lobes spreading, narrowly elliptic 4.5 – 5 x 2 – 2.5 mm, reflexed along the midline, marginal 0.3 – 0.4 mm hyaline, white, undulate, lacking trichomes, central part densely covered in papillae, glabrous, outer perianth 10 – 20% curved in purple-black stellate hairs (see pedicel). Androphore cylindrical, or narrower at apex than base, (1 –)1.3 x 0.4(– 0.6) mm, c. 10% covered in 5 – 8-armed dark brown stellate hairs 0.1 – 0.25 mm diam (Fig. 3F); anther-head disc-like, 0.7 x 1.4 – 1.5 mm, glabrous; stamens 7 – 8, anthers narrowly elliptic, 0.6 x 0.3 – 0.4 mm. *Gynoecium vestigial* 0.3 mm diam., 0.15 mm long, carpels 5, each asymmetrically ovoid, laterally flattened, with minute, sparse stellate hairs; style-stigmas apical, conical, erect. *Female flowers and Fruits* unknown.

**DISTRIBUTION & HABITAT.** Congo (Brazzaville), Massif du Chaillu, evergreen forest with Detaroid legume canopy, nearly submontane; 640 – 660 m alt.

**ADDITIONAL SPECIMEN.** Republic of Congo (Congo-Brazzaville), Lekoumou Prefecture, N of Lewalla, plot 33, 2°41’ 03” S, 13°35’ 29” E, st. 21 May 2010, *E. Kami* 4719 (IEC!, K!).

**CONSERVATION STATUS.** Known from a single plant with certainty, however, a second sterile specimen from a plot voucher at a second location is attributed to this species (see additional specimen). This species is extremely rare since although many thousands of specimens were collected at and around the Zanaga project location, only two were collected of the new species. The species should be assessed as Endangered since there is uncertainty as to future management of the locations and there are threats of mining and logging.

**PHENOLOGY.** Flowering probably in late wet season - early dry season (collected in late dry season with dead flowers).

**ETYMOLOGY.** Named for the nearest large town, Zanaga.

**LOCAL NAMES & USES.** None are known.

**NOTES.** collected in 2009, *Cheek* 15853 was initially identified as the closely similar and Endangered *Cola mayimbensis* the type collection of which was made in adjoining Gabon, at Lastoursville. The taxonomic status was re-assessed after Y.B. Harvey (WRSY) pointed out that the leaf-blades of *C. mayimbensis* are half the size of those of *Cola zanaga*. This observation promoted more critical scrutiny during which numerous additional points of difference between the taxa were recorded (Table 2).

**Table 2.** Characters separating *Cola mayimbensis* from *Cola zanaga.* Characters for *Cola mayimbensis* from Hallé (1961) and from observations of material cited at P.

|  | *Cola mayimbensis* | *Cola zanaga* |
| --- | --- | --- |
| Leaf-blade shape | Elliptic | Oblanceolate (rarely obovate-elliptic) |
| Leaf-blade dimensions (cm) | 5.5 – 12 x 1.7 – 4 | (10 – )11.7 – 17.8 x (4 – )4.7 – 6( – 6.2) |
| Petiole length (mm) | 4 – 6 | (2.5 – )5 – 10 |
| Androphore length (mm) | 2 | 1 – 1.3 |
| Petiole and stem indumentum | Densely simple hairy (100% cover), persistent | Sparsely (only 10 – 20% cover) hairy, caducous |
| Stipules caducity, indumentum | Extremely caducous, rapidly glabrescent | Persistent, densely & persistently hairy |
| Perianth shape, posture | Campanulate, pendant | Lobes spreading, erect |
| Perianth lobe: tube ratio | 1:1 | 5:1 |
| Geography | Gabon | Congo |

Additional new species recorded from this site in the Massif du Chaillu of Congo (Brazzaville) are *Octoknema chailluensis* (Gosline & Malécot 2012) and *Gilbertiodendron quinquejugum* Burgt (van der Burgt *et al*. 2015). Republic of Congo as a whole continues to yield new discoveries of threatened plant species (e.g. Gosline *et al*. 2014; Langat *et al*. 2021).

The absence of recorded female flowers from the single specimen suggests that the species may be dioecious. However, the paucity of flowers on the specimen does not make this conclusive, and monoecy cannot be completely ruled out.

#### 10. Cola moussavoui

Breteler (2014a: 102). Type: Gabon, Ngounié, Bindolo R. basin, NW of Fougamou, fl. 20 Sept. 1997, *Breteler, Leal, Moussavou & Nang* 14011 (holotype WAG0091301; isotypes WAG0029782, WAG0091302).

**DISTRIBUTION & HABITAT.** Gabon, known only from the type location; lowland evergreen forest.

**ADDITIONAL SPECIMENS.** None are known.

**CONSERVATION STATUS.** *Cola moussavoui* was assessed as Critically Endangered (CR B1ab(iii)+2ab (iii)) by Cheek & Baldwin (2019b). The single location, where it is known from one collection, is threatened with ongoing extensive slash and burn agriculture and clearance for plantations (Cheek & Baldwin, 2019b).

**PHENOLOGY.** Flowering in Sept., fruiting unknown.

**ETYMOLOGY.** Named for Jean-Mathieu Moussavou, field botanist and technician of the Herbier National du Gabon (LBV), one of the collectors of the type material (Breteler 2014 105).

**LOCAL NAMES & USES.** None is known.

**NOTES.** *Cola moussavoui* is most closely similar morphologically to another Gabonese endemic, *Cola mayimbensis* Pellegr. Remarkably, Breteler did not compare the two species. Both species share elliptic leaf-blades of similar dimensions and ratios, with narrowly triangular acumen, 5 – 7 lateral nerves on each side of the midrib, petioles c. 5 mm long, male flowers with 5 tepal lobes, each with white membranous, undulate fringes and a length of 2.5 – 3 mm. However, the pedicels of *C. mayimbensis* can be up to twice as long as those of *C. moussavoui,* the androphore is more than twice as long, and bears 8 not 5 anthers, while the stipules, which are subulate, with a length:breadth ratio of 6:1 and have a raised midrib, are quite different to those of *C. moussavoui* which are lanceolate, with a ratio of c. 3:1 and lack a raised midrib. However, these may well be sister species.

There is no indication that more than a single plant was found, and this is supported by just three duplicates being indicated. If this is the case, since both male and female flowers are described, it seems likely that *Cola moussavoui* is monoecious.

It was not possible to view images of the three type sheets. As with *Cola stigmatosa* (above) this species appears not yet to have been register on the Naturalis website (18 July 2022), however the illustration in the protologue is of excellent quality.

New species to science, often highly range-restricted, continue to be discovered in Gabon e.g. eight new species of *Palisota* (Bidault & van der Burg 2019), six new species of *Monanthotaxis* (Hoekstra *et al*. 2021), numerous *Psychotria* (Lachenaud 2019). However, some appear extinct even before they are published such as *Pseudohydrosme bogneri* Cheek & Moxon-Holt (Moxon-Holt & Cheek 2021). In 2019 alone, 37 new species to science were published from Gabon, second only to Cameroon of all tropical African countries (Cheek *et al*. 2020b).

#### 11. Cola toyota

Cheek sp. nov. Type: Cameroon, Centre Region “Forêt sommitale du Mont Meza-feuille IGN-1/200.000 NANGA EBOKO”, 900 – 1000 m alt., fl.fr., 12 May 1959, *Letouzey* 1956 (holotype YA!; isotypes K barcode K000874907!, P!) (Fig. 4).

**Fig. 4.**
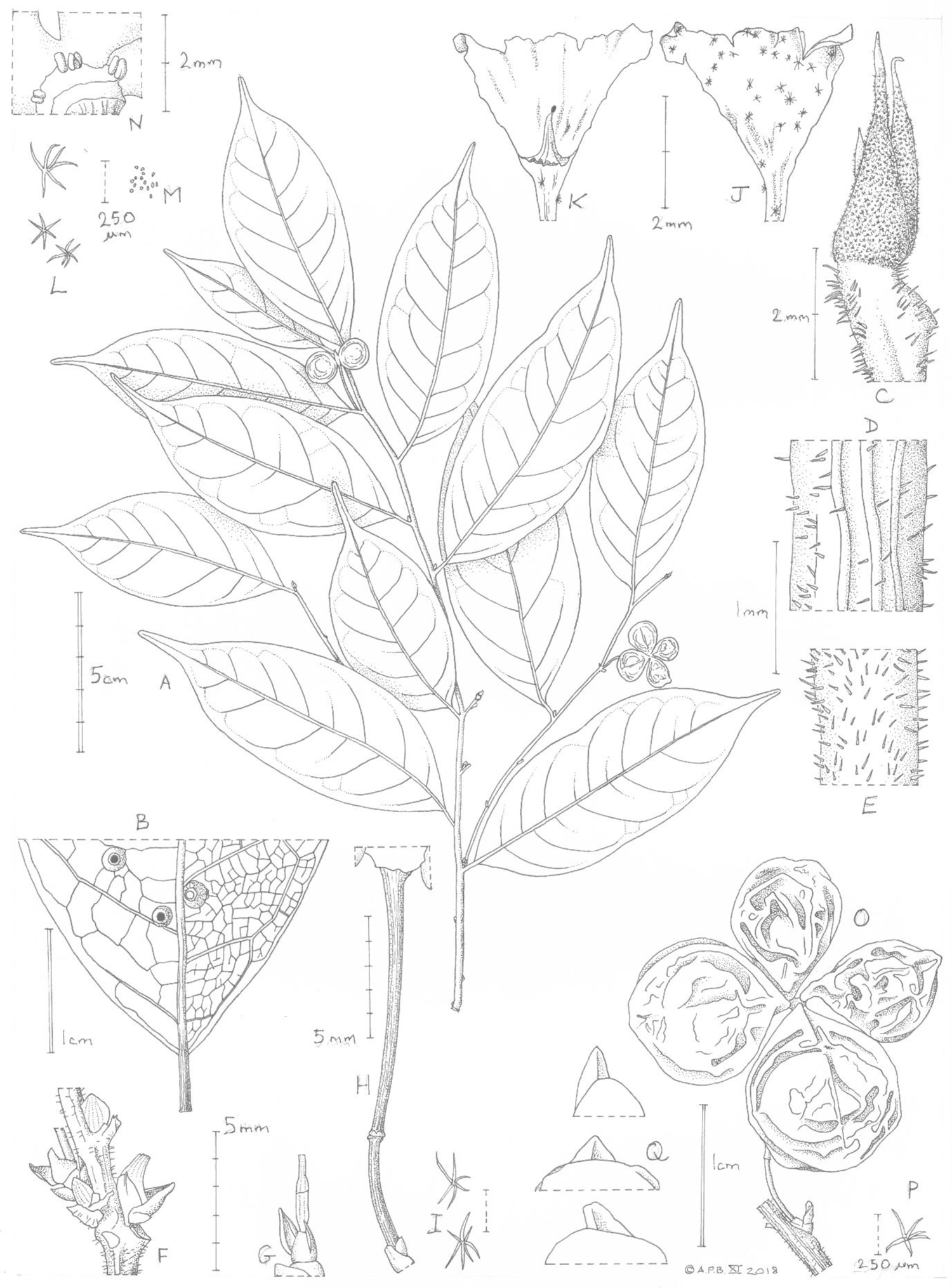
*Cola toyota.* **A.** habit, fruiting, leafy stem; **B.** detail of abaxial leaf surface showing galls; **C.** stem apex and terminal bud with stipules; **D.** stem surface and indumentum; **E.** petiole indumentum; **F.** detail of inflorescence bract; **G.** base of inflorescence showing articulated pedicel; **H.** base of fruiting (accrescent) pedicel; **I.** stellate hairs from pedicel; **J.** side view of old male flower (incomplete); **K.** inner surface of J showing androphore; **L.** stellate hairs from outer surface of perianth in J; **M.** papillae from inner surface of perianth in **K**; **N.** androecium – persistent stamens at base of fruiting carpels; **O.** fruit; **P.** stellate hair from fruit surface; **Q.** examples of fruitlet rostra. All from *Letouzey* 1956. By Andrew Brown. nerves conspicuous on abaxial, not adaxial surface, slightly raised on the lower surface, white, forming a reticulum with isodiametric cells, glabrous. *Petioles* terete, (3.5 –)4 – 6.5 x 0.8 mm long, colour and indumentum as stem, but hairs denser c. 30% cover (Fig. 4 E); pulvini absent. Stipules caducous, not seen. *Inflorescences* only known from old flowers and from remains with fruits, axillary, single or few-flowered fascicles (Fig. 4 F&G), 1 inflorescence per leafy stem, at about the third node from the stem apex; peduncle not seen; pedicels 2.5 – 4.5(– 9) mm long below the articulation, dilated (thicker) at the articulation (Fig. 4 G&H), 11 – 17 mm long above the articulation (15 – 26 mm long in total), sparsely stellate hairy, hairs colourless, 6 – 7 – armed, arms spreading, 0.25 – 0.4 mm diam. (Fig. 4 I).*Galls* occasional, on lower surface of blade, volcano-like, 1.5 – 2 mm diam., the central crater 1/3 – 1/2 the total diameter (Fig. 4 B). *Bracts* 3(– 5), clustered at base of inflorescence, brown, indurated, glabrous ovate, the outermost (proximal) flat, c. 0.5 mm long, the more distal bracts longer, 1 – 2 mm long, the most distal naviculate (boat-like), sometimes acuminate, or the whole bract subulate-naviculate, sometimes longitudinally bifurcate (Fig. 4 F,G&H). *Male flowers* (old, incomplete) colour unknown. *Perianth* tube funnel-shaped 2.5 – 3 x 3.5 mm distal part divided into 5 lobes, lobes (incomplete, damaged) each 1 mm wide at base (Fig. 4 J&K), lacking wide membranous, undulate margins; outer surface of funnel and lobes with sparse 5 – 8-armed stellate hairs covering 5 – 10% of surface (at least the proximal part of the lobes, distal part not present), hairs 0.2 – 0.4 mm diam. (Fig 4 L); inner surface with minute translucent vesicles 0.02 mm diam. (Fig. 4 M). Androphore 0.75 x 0.1 mm long, tapering from a broad, fluted base to a narrow apex, glabrous. Anther head not preserved. *Female flowers* (remnants observed in fruit), with anthers 5 (Fig. 4 N), ellipsoid 0.2 – 0.5 mm long, glabrous. *Fruit* with 4 or 5 carpels, gynophore 2 x 1.5 mm, carpels patent, globose, 0.8 – 1.2 c diam., stipe inconspicuous, apex shortly mucronate-rostrate, the beak triangular in side view, 0.2 x 0.2 cm (Fig.4 G). Surface glossy, drying dull yellow, with scattered stellate hairs, diam 0.8 – 1.2 mm, arms 5 – 7, spreading. *Seeds* not investigated due to paucity of material.

*Monoecious shrub,* probably several metres tall, trunk 10 cm diam, straight, greenish, with lenticels, bark detaching in strips, slash yellow. *Leafy stems* terete, c. 2 mm diam. at lowest leafy node, dark purple-brown, longitudinally ridged, lenticels dense elliptic, raised, bright white; glabrescent, sparsely hairy at stem apex, 10 – 20% covered in erect, simple, colourless hairs c. (0.05 –)0.1(– 0.15) mm long (Fig. 4 D); internodes 18 – 27 mm long; scale-leaves absent. *Terminal and axillary buds* acute, 3.5 – 4 x 1.2 mm, stipules loosely appressed to bud, densely (c. 90% covered) minutely stellate-hairy, hairs 3 – 5-armed, 0.05 – 0.07 mm diam. (Fig. 4 C). *Leaves* 3 – 10 over the length of a season’s growth, persisting for two seasons or more, drying black on upper surface, slightly asymmetrically elliptic, rarely elliptic-lanceolate, (6 –)8.8 – 11.3 x (2 –)3 – 3.9(– 4.2) cm, blade 0 – 7 mm wider on one side of the midrib versus the other (Fig. 4 A), acumen straight or curved, (0.8 –)1.3 – 1.7(– 1.9) x (0.2 –)0.3 – 0.4(– 0.5) cm, apex rounded. Midrib raised, white, secondary nerves (7 –)8 – 9 on each side of the midrib, arising at 50 – 70° from the midrib, towards the margin arching upwards towards the apex and uniting with the secondary nerve above, forming a looping infra-marginal nerve 2 – 5.5 mm from the margin, domatia absent; tertiary and quaternary

**RECOGNITION**: differs from *Cola mayimbensis* Pellgr. in the stipules highly caducous, densely stellate hairy (vs persistent for a whole season, simple hairy), fruit apex shortly rostrate (vs rounded); perianth lobe margins thickened, (vs broad, membranous, white and undulate).

**DISTRIBUTION & HABITAT**”assez abondant en sous bois de forêt de type primaire à *Celtis mildbraedii* (altitude 900 – 1000 m).” (Common in the understory of primary *Celtis mildbraedii* forest 900 – 1000 m alt.). Collected with (*Letouzey* 1955 (YA), same location & date) the type of *Ardisia dolichocalyx* Taton (Primulaceae, Taton 1979), also a forest understorey shrub species.

**ADDITIONAL SPECIMENS.** None are known

**CONSERVATION STATUS.** *Cola toyota*, is known from a single location, Mount Meza, where it is threatened by clearance of its forest habitat for agriculture. Viewing historical imagery of Mount Meza on Google Earth Pro (accessed 4 May 2020) it is possible to see that forest clearance by 2014 had advanced from the West and North to the type locality at the summit of Mt Meza. While forest still existed in 2014 (latest available imagery) in the vicinity of the summit, it is possible that clearance is still extending and reducing the habitat of this species. Therefore, *Cola toyota* is here assessed as Critically Endangered, CR B1ab(iii) +B2ab(iii) since area of occupancy is 4 km² using the IUCN (2012) preferred cell-size of that dimension, and extent of occurrence is taken as equal. It is to be hoped that the species will be found at other locations, but it is possible that it is unique to this isolated mountain top as are other species of *Cola* to other mountains, e.g., *Cola etugei* Cheek, unique to Mt Kupe in SW Region (Cheek *et al*. 2020a). Although *Cola toyota* has not been recorded for 60 years, this is probably because no botanist has revisited the location.

The georeference for the type locality is deduced to be 4°16’02.71” N, 12°07’19.28” E. This point, read from Google Earth Pro (accessed 4 May 2020) gives the peak of a mountain (summit reading 978 m) nearest (1.2 km) to the site labelled (on Google Earth Pro) as the village of Meza, 50 km SW of Nanga Eboko.

**PHENOLOGY.** Flowering and fruiting in May.

**ETYMOLOGY.** Named for the Toyota Motor Corporation for their part in supporting *Cola* extinction risk assessments at RBG, Kew (see acknowledgements).

**LOCAL NAMES & USES.** None recorded.

**NOTES.** *Letouzey* 1956, the sole specimen known of *Cola toyota*, bears fruits and a few old and incomplete male flowers as well as fruits, so the species is monecious. The initial determination by Letouzey was “*Cola cf. mayimbensis* Pellegr.”, but his final determination made at Paris was “*Cola philipi-jonesii* Brenan & Keay”, dated May 1964. This may indicate that Letouzey compared his specimen with the type of *C. mayimbensis* at P and found that it did not match, concluding that it must be *C. philipi-jonesii*, which is not represented at P.

The sheet at Kew is later annotated in the hand of Verdcourt “*Cola cf. mayimbensis* Pellegr. (leaf apex of *philipi-jonesii* is different and indumentum)”. This indicates that Verdcourt, who had access to the type of *Cola philipi-jonesii*, considered the specimen near to but different from *C. mayimbensis.* In fact, the purple-brown epidermis of the stem with white, raised lenticels remarked on by Letouzey are shared with *Cola mayimbensis*, not *Cola philipi-jonesii*, and *Cola mayimbensis* also shares with the *Cola toyota* the patent simple translucent indumentum of the stem, although this is denser and more persistent in *Cola mayimbensis* (in *Cola philipi-jonesii* the indumentum is mainly stellate, not simple). *Cola toyota* shares several other features with *Cola mayimbensis*. Since the flowers of the first are very incompletely known, it is difficult to compare them on that basis. However, the two can be separated using the features shown in Table 3 and in the diagnosis. Both species occur in submontane habitats but are separated by about 360 km of lowland evergreen forest.

**Table 3.** Diagnostic characters separating *Cola mayimbensis* from *Cola toyota*. Data for the first species taken partly from Hallé (1961:66 – 68).

|  | *Cola mayimbensis* | *Cola toyota* |
| --- | --- | --- |
| Fruit apex | Rounded | Rostrate |
| Stem indumentum | Densely (80% cover), persistently hairy | Sparsely hairy (10-20% cover) glabrescent |
| Inflorescences frequency per leafy stem | Numerous, in successive axils of a stem | One fertile node per stem |
| Stipule indumentum | Hairs simple | Hairs stellate |
| Stipule persistence | Persistent for the entire season's growth | Caducous, seen at apical bud only |
| Perianth lobe margins | With a wide, membranous, undulate band | Margin thickened, not membranous or undulate |
| Leaf-blade galls | Absent | Volcano-like galls conspicuous on lower blade surface |
| N°. secondary nerves on each side of the midrib | 5 – 7 | (7 – )8 – 9 |
| Reticulate 3° & 4° nervation conspicuous on adaxial blade surface: | Yes | No |
| Number of anthers | 8 | 5 |

The habitat of *Cola toyota* is likely to be submontane forest. Although the canopy tree is given as a *Celtis*, indicative of semi-deciduous forest, the presence of *Cola* and *Ardisia* in the understorey, at 900 – 1000 m alt. where cloud interception is usual, is indicative of submontane or cloud forest, usual at above 800 m alt. in Cameroon. (Cheek pers. obs.

Cameroon 1984 – 2016). This vegetation has been found to be rich in narrow endemics e.g., *Coffea montekupensis* Stoff. and several species of *Kupeantha* (Rubiaceae, Stoffelen *et al*. 2007; Cheek *et al*. 2018d) where it survives and has been studied in SW and NW Regions, Cameroon (e.g. Cheek & Onana 2021; Cheek *et al*. 2022c). However, that of the Nanga Eboko area seems almost completely unstudied apart from Letouzey’s visit in 1959. Further survey effort is called for to elucidate the species present, in the expectation that additional new species to science are likely to result.

#### 12. Cola takamanda

Cheek sp.nov. Type: Cameroon, South West Region, Takamanda Forest Reserve, near Matene. 6°14’N, 9°19’E, 170 m alt. fl. 21 – 25 March 1985, *D. Thomas* 4536 (holotype YA! isotype MO n.v.) (Fig. 5).

**Fig. 5.**
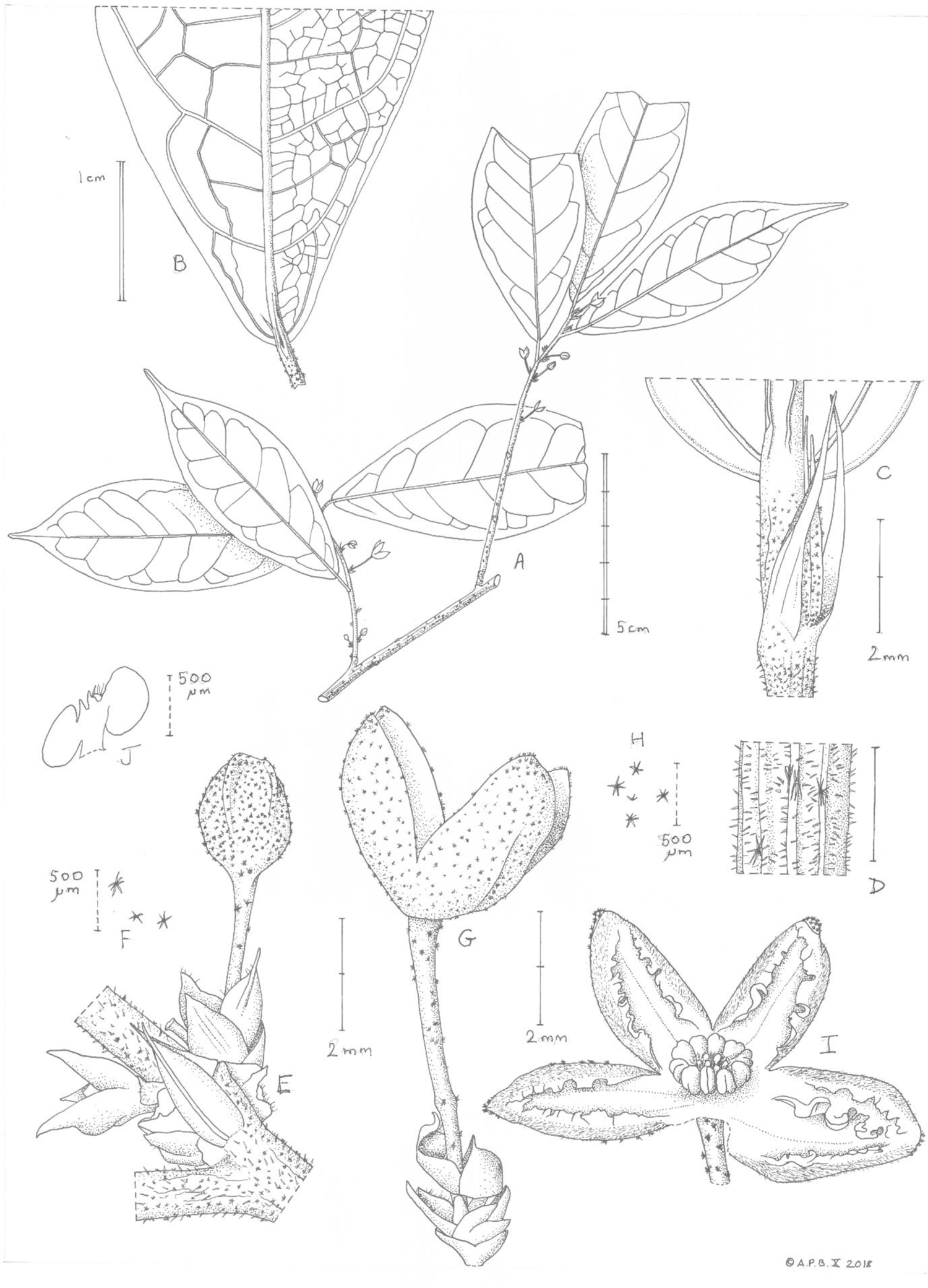
*Cola takamanda.* **A.** habit, flowering, leafy stem; **B.** abaxial leaf surface showing galls; **C.** leaf base and stipules at stem apex; **D.** stem surface and indumentum; **E.** inflorescences, with a flower bud; **F** stellate hairs from pedicel; **G.** inflorescence with flower and bracts; **H** stellate hairs from abaxial tepal; **I** male flower; **J** section through androecium showing short androphore. All drawn from *Thomas* 4536 (YA) by ANDREW BROWN.

*Dioecious or monoecious, probably evergreen, small tree,* height and architecture unknown. *Bud-scales* not seen. Internodes more or less even in length, 3.5 – 7.5 mm long. Stem of current season’s growth dark purple, 0.8 – 1.5 mm diam. with fine longitudinal ridges, 30 – 40% covered in hairs of two types, 1) predominantly patent translucent simple hairs 0.05 – 0.1 mm long; 2) c. 5% cover of stellate 5 – 7 – armed hairs 0.25 – 0.3 mm diam., arms directed mainly to base and apex of stem axis (Fig. 5C & D). Stem of previous season’s growth with raised circular white lenticels 0.8 – 1.1 mm diam., sometimes with a central longitudinal groove, simple hairs mostly persistent, stellate hairs caducous. *Leaves* – 3 per season’s growth, alternate, distichous, more or less uniform in size and shape throughout the season. Leaf-blades concolorous, grey-green narrowly elliptic or slightly oblanceolate 8 – 10.5 x 2.5 – 3.5(– 4) cm, acumen narrowly triangular 0.75 – 1.8 cm long, apex acute, finally abruptly rounded, base cuneate, finally abruptly rounded to truncate, secondary nerves 6 (– 7) on each side, arising at 50 – 70° from the midrib, curving upwards steeply about 2/3 the distance to the margin and uniting with the nerve above (brochidodromous), forming a conspicuous looping infra-marginal nerve 1 – 9 mm from the margin; tertiary nerves uniting with the intersecondary nerve to form a zig-zag line (Fig. 5B); tertiary nerves conspicuous, reticulate, cells oblong c. 1 – 1.2 mm long and uniting outside the marginal looping infra-secondary nerve to form a second, finer, parallel infra-marginal nerve c. 0.5 mm from the margin. *Galls* present, on adaxial surface of leaf initially a glossy brown cone 0.3 mm diam., the surrounding cuticle later lifting, forming a low dome 1.5 mm diam., glabrous. The corresponding part of the lower surface with dome 1.5 – 1.7 mm diam., hairs long, sparse, simple translucent to pale brown c. 1 mm long. *Petioles* terete, (2.5 –)3 – 5.5 x 0.8 – 1 mm, lacking pulvini, c. 50% covered in translucent simple patent hairs c. 0.1 mm long mixed with a few 4 – 6-armed stellate hairs 0.1 – 0.15 mm diam. *Stipules* subulate, 4.2 – 4.5 x 0.75 mm, glabrous, indurate, brown. *Inflorescences* axillary, 1-flowered or fasciculate on current season’s growth, at 5 – 7 nodes (with or without leaves) per stem; 1 – 2(– 3)-flowered, flowers opening in succession. *Bracts* 4 – 5, alternate, spirally arranged, proximal (basal)-most bracts deeply bifurcate, the two lobes each lanceolate, 1 – 1.8 x 0.2 – 0.7 mm; middle bracts lanceolate 1.5 – 2.7 x 0.5 – 0.8 mm, apex with an awn 0.4 – 0.6 mm long, extending from the conspicuously raised midrib, margin with spreading simple hairs 0.1 mm long; distalmost bracts hooded – naviculate, 1.2 – 1.5 mm long, apex beaked, glabrous. *Pedicel* 4 – 7 mm x 0.3 mm, widening to 0.5 mm diam. at apex, articulation concealed (by bracts), c. 1 mm from base, indumentum c. 5% cover, of 5 – 7-armed stellate hairs 0.15 – 0.2 mm diam. *Male flowers* yellow-brown. *Perianth* divided by 8/10 into (4 –)5 patent lobes, tube bowl-shaped, 1.5 x 3.5 – 4 mm; lobes each 2.8 – 3 x (1.3 –)1.5(– 2) mm (Fig.5I), the margins wide, membranous, undulate and inflexed, 0.5(– 1 mm wide), outer surface glabrous in the marginal half, the remainder densely covered in appressed simple hairs; outer surface of perianth 20 – 30% covered in (3 –)6 – 7-armed orange-red stellate hairs (0.06 –)0.12 – 0.17 mm; inner surface with minute translucent vesicles 0.02 mm diam., otherwise glabrous. Androphore terete, 0.1 – 0.2 mm long, glabrous; anthers five, forming a head 1.5 mm diam., anther cells elliptic, 0.5 x 0.25 – 0.3 mm; vestigial carpels six, subcapitate, glabrous, surrounded by simple hairs and concealed within the staminal ring. *Female flowers* and *Fruit* unknown.

**RECOGNITION.** Differing from all other species of *Cola* in the extremely contracted androphore of the male flowers (0.1 – 0.2 mm long, vs >0.75 mm long), further differing from *Cola philipi-jonesii* Brenan & Keay of Nigeria in the narrowly triangular acumen with apex acute (vs spatulate); perianth lobes with a broad, inflexed membranous margin (vs with margin undifferentiated). Further differing from *Cola mayimbensis* Pellegr. of Gabon in that the carpels of male flowers are concealed within the staminal ring (vs the carpels exserted from the staminal ring and plainly visible), the stipules glabrous, (vs densely simple hairy).

**DISTRIBUTION & HABITAT.** Cameroon: South West Region, Takamanda Forest Reserve; “mature rainforest with *Cynometra hankei, Terminalia ivorensis, Strombosia glaucescens* and *Anonidium* sp.” (*Thomas* 4356, YA)

**ADDITIONAL SPECIMENS.** None are known.

**CONSERVATION.** *Cola takamanda* is known from a single collection at one site. According to the specimen grid reference, this was within the Takamanda Forest Reserve, which since 2008 has been named the Takamanda National Park. However, the grid reference is given as degrees and whole minutes on the label, confirming the view that a Global Position System (GPS) was not used for the collection in 1985. GPS were generally not used in botanical surveys until the 1990s. Therefore, the grid reference is likely to have been read from a map and so be imprecise. The label also states ‘Near Matene’ which is outside the National Park. Therefore, it is possible that the record is on the edge of, or outside of the National Park boundary where it is at threat from slash and burn agriculture prevalent in the area (observed on Google Earth imagery). For this reason, using the precautionary principle, we here assess *Cola takamanda* as Critically Endangered (CR B1+B2ab(iii)).

It is quite possible that, as with similar and geographically close short-petioled *Cola* species, such as *Cola philip-jonesii* (also known from a single site) that this species is locally common within its sole global locality. While *Cola takamanda* may be more widespread, it has not been found during surveys of other forest areas in western Cameroon and adjoining areas (Cheek 1992; Cheek *et al*. 1996; Cable & Cheek 1998; Cheek *et al*. 2000; Maisels *et al*. 2000; Chapman & Chapman 2001; Cheek *et al*. 2004; Harvey *et al*. 2004; Cheek *et al*. 2006; Cheek *et al*. 2010; Harvey *et al*. 2010; Cheek *et al*. 2011). Therefore, it may truly be endemic to Takamanda. This mirrors the situation with *Begonia stellata* Sosef (Sosef 1994), also known from a single collection by Duncan Thomas at Takamanda and assessed as Critically Endangered (Onana & Cheek 2011:103). Takamanda and the adjoining lowland areas of Cameroon to the East are extremely incompletely (or even not at all) surveyed for plants, but where surveys have been mounted, other similar narrow endemic species are recorded, e.g. *Psychotria monensis* Cheek & Séné from the nearby Mone Forest Reserve (Sené & Cheek 2010), *Memecylon bakossiense* R.D.Stone *et al*. (2008), or *Warneckea ngutiensis* R.D. Stone at Nguti (Stone & Cheek 2018). We recommend that *Cola takamanda* be incorporated within the management plan of the Takamanda National Park and that the population be surveyed, mapped and monitored for survival. Since *Cola* species only grow in good quality, intact forest, it is recommended that efforts be made to work with the local community in Matene to avoid slash and burn agriculture in the zone where *Cola takamanda* occurs and to provide training on identifying the species and understanding its global importance for conservation. The Takamanda Forest Reserve, established in 1934, is best known in the conservation world as one of the few areas in which the Cross River Gorilla, *Gorilla gorilla diehlii* survives. This is the most western and the most northern gorilla taxon, and also the world’s rarest great ape, since only c. 300 individuals are thought to survive in 10 – 14 locations (Sarmiento & Oates, 2000).

**PHENOLOGY.** Flowering in March.

**ETYMOLOGY.** Named for the forest reserve (and village) of Takamanda, to which this species is unique, on current evidence.

**NOTES.** The type and only known specimen of *Cola takamanda* was annotated as *Cola mayimbensis* by Nkongmeneck in 1988 (YA). It is indeed similar to that species in general aspect, in the shape of the leaves, the broad, inflexed, undulate, perianth margin but differs in the features indicated in Table 3 and in the diagnosis.

The most remarkable single feature of *Cola takamanda* is the contracted androphore of the male flowers which is far shorter than the length of the anther cells in the anther ring which it supports. In all other species of the genus, the androphore of the male flowers is far longer than the length of the anther ring.

The absence of female flowers from the single specimen suggests that the species may be dioecious. However, the paucity of flowers on the specimen does not make this conclusive, and monoecy cannot be completely ruled out.

### Incompletely known species

The following species appear to be distinct from all other known species of subg. *Disticha* based on their morphology, however no flowering or fruiting material is available. The current taxonomic convention is that such species should not be formally named until reproductive material is available.

**A. Cola ‘Campo-Ma’an** Specimen: Cameroon, South Region Ebianemeyong, Campo-Ma’an N. Park, st., date unknown (probably c. 2000), *Tchoutou* EBOX 181 (K!).

*Shrub* height unknown. Stem black, densely (c. 80% cover) simple-hairy, hairs persistent for 5 – 9 internodes from the apex, hairs 0.15(– 0.4) mm long, erect, translucent. Internodes 0.6 – 1.6(– 1.9) cm long. Older internodes (distant from apex), with bright white raised orbicular lenticels 0.5 mm diam. *Terminal buds* loose, open, ellipsoid c. 3.5 x 1.9 mm, with 9 – 10 scales; bud-scales narrowly triangular to naviculate, brown, indurated, glabrous 2.8 x 0.6 mm, midrib raised. *Leaves* elliptic 8.8 – 10.3 x (2.8 –)3.2 – 4(– 5) cm, acumen 1.7 – 1.8 cm long, 0.3 cm wide, apex rounded, drying pale green, nerves white. Secondary nerves 6 – 7 on each side of the midrib, arising at c. 50° from the midrib, curving upwards and uniting with the nerve above, forming a looping nerve 3 – 4 mm from the margin; tertiary and quaternary nerves forming a conspicuous raised reticulum on both adaxial and abaxial surfaces. *Leaf galls* absent. *Petioles* black, indumentum as stem, ± terete 4 – 6 mm long. *Stipules* caducous, only at stem apex and second node, narrowly triangular-subulate, 3.8 – 5 mm long, 0.5 mm wide, midrib raised, glabrous, apart from a few stellate hairs at base. *Flowers and Fruit* unknown.

**DISTRIBUTION & HABITAT.** CAMEROON, South Region, Campo-Ma’an National Park. Lowland evergreen forest.

**CONSERVATION STATUS.** *Cola* ‘campo-ma’an’ would likely be assessed as Critically Endangered under criterion B of IUCN (2012), since there are clear threats in its single location.

**NOTES.** the sole specimen known was donated by J.J.F.E. de Wilde at WAG to K for confirmation of the identification. The specimen is a plot voucher collected as part of Peguy Tchouto’s doctoral research (Tchouto 2004), conducted at WAG, on the plants of Campo-Ma’an. The specimen had been identified as *Cola philipi-jonesii* by Tchoutou in Oct. 2000, confirmed by Breteler in April 2002. This was a reasonable determination at the time since only two species of the subgenus were then known from the whole of the Lower Guinea phytogeographical zone, namely the species indicated and *C. mayimbensis* of Gabon. With the benefit of data available today, we can rule out *C. philipi-jonesii* because that species has dense stellate (not simple) indumentum that sheds to reveal a grey-white (not black) epidermis, and the acumen is spatulate, not narrowly oblong, as wide at apex as at base.

Tchoutou’s specimen differs from *C. mayimbensis* because that species has persistent, densely simple hairy stipules (vs caducous, ± glabrous) and leaves with the quaternary and tertiary nerves inconspicuous and not raised on the adaxial surface. Initially this specimen was aligned with the species described in this paper as *Cola toyota*. However, in that species the stem is only 10 – 20% covered in indumentum (vs c. 80%) and the stipules are densely stellate hairy (not glabrous or with just a few stellate hairs) and also the tertiary and quaternary nerves are inconspicuous on the adaxial surface (vs raised and conspicuous).

Finally, in *C. toyota* conspicuous galls are present on most leaves, yet they are absent in the *Tchoutou* specimen. There seems no doubt that this is a distinct and unnamed species of *Cola* subg. *Disticha*.

**B. *Cola ‘udzungwa*’** Specimen: Udzungwa Mts National Park, 1330 m alt. st. 30 Oct. 2005, *Luke WRQ, Mwanoka M, Festo L* 11311 Tanzania, (EA!, NHT).

*Tree* 3 m. Stem sparsely (c. 5% cover) patent, simple hairy, hairs translucent, 0.08 mm long, persisting for 2 – 3 nodes, distal 2 – 4 nodes green, more proximal nodes white-pale brown, longitudinally ridged and furrowed, lenticels ± concolorous, dull white orbicular, 0.5 mm diam., internodes (1.3 –)1.9 – 2.2 cm long. *Terminal buds* not seen. *Leaves* oblong-elliptic, (10.2 –)12.3 – 15 x (3.4 –)4 – 4.6 cm, acumen twisted, narrowly triangular, 1.1 – 1.8 x 0.2 – 0.4 mm, base rounded or obtuse, drying pale green, veins dull white. Secondary nerves 8(– 9) on each side of the midrib, arising at 50 – 60° from the midrib, arching upwards and forming a poorly defined, incomplete nerve 3 – 4 mm inside the margin; tertiary and quaternary nerves forming a reticulation conspicuous in the lower surface, but not on the upper, adaxial surface; lower surface with scattered minute sessile red globose glands c. 0.025 mm diam. *Galls* absent. *Petioles* yellow-green (5 –)7(– 8) mm long, indumentum densely (90 – 100% covered) hairy, hairs pale brown stellate 0.1 – 0.3 mm diam., 5 – 9-armed, arms appressed, mixed with minute erect simple hairs. *Stipules* caducous, not seen. *Flowers and fruits* unknown.

**DISTRIBUTION & HABITAT.** TANZANIA, Udzungwa Mountains; submontane forest on steep slope, 1330 m alt.

**ADDITIONAL SPECIMENS. TANZANIA.** Udzungwa Mts National Park; submontane forest 900 m alt., Nr. Camp 244, *Luke PA & Luke WRQ* 8747 (EA!, K!).

**CONSERVATION STATUS.** *Cola* ‘udzungwa’ cannot be assessed for its conservation (extinction risk) assessment because it cannot be formally named: fertile material is lacking. However, were this obstacle overcome, it is likely that with a single ‘threat-based’ location, and threats, it would likely be assessed as Critically Endangered under criterion B of IUCN (2012), if threats can be adduced.

**NOTES.** Although sterile, this taxon, although tentatively identified as *Cola uloloma* by Quentin Luke and despite the similarity to that species, and the lack of reproductive material, is clearly a different species. It lacks the white-waxy stem coating of *Cola uloloma*, while the simple hairy stems, densely stellate hairy petioles, rounded leaf-bases, high altitude cloud forest habitat, and red glandular abaxial leaf-blade contrast with the glabrous stems, sparsely stellate petioles, cuneate leaf-bases, low altitude semi-deciduous forest habitat, and non-glandular abaxial leaf-blades of *C. uloloma*.

*Luke & Luke* 8747 also from Udzungwa, 900 m alt., has oblanceolate leaf-blades unlike those of *Luke et al.* 11311, and has simple hairy petioles rather than stellate and simple hairy petioles. In all other respects, it matches *Luke et al.* 11311. The differences in leaf blade shape may be due to being from a juvenile versus a mature plant, and the difference in indumentum due to the great caducity of stellate versus simple hairs. A mission to the Udzungwa’s, targeting to find this species in flower or fruit, so that it might be more fully characterised and formally named is recommended.

New narrowly endemic and threatened species to science are regularly being described from Tanzania (e.g. Champluvier & Darbyshire 2009; Cheek & Bridson 2019; Johnson *et al*. 2017), including also from the Udzungwa Mts e.g. *Vepris udzungwa* Cheek and *V. lukei* Cheek (Cheek & Luke 2022)

## Acknowledgements

The stimulus to reawaken the dormant monographic study of *Cola* was provided by sponsorship via IUCN from the Toyota Motor Corporation to the RBG, Kew to establish a Plant Assessment Unit and increase the rate of completion of extinction risk assessments of tropical plant species for publication on the IUCN Red List. Special thanks to Steve Hope. Eimear Nic Lughadha (PI) and Serene Hargreaves (Project Manager) prioritised conservation status assessment of *Cola* species, following the completion of the global assessment of wild coffee species. Poppy Lawrence and Isabel Baldwin are thanked for their research on the extinction risk assessments of those *Cola* species in this paper that had been previously published. Recently the above target has been combined with the need for data to define Important Plant Areas (IPAs or TIPAs) in Cameroon necessitating renewed work on species delimitation of that genus by the author to support these. The Cameroon TIPAs (Tropical Important Plant Areas) project is supported by Players of Peoples Postcode Lottery. We thank Lydia Burns of Kew’s Foundation for facilitating this. Gaston Achoundong, former head of the National Herbarium of Cameroon (YA) and his successors especially Jean Michel Onana are thanked for their collaboration and support hosting visits to study *Cola* specimens over the years. The curators of several other herbaria, BM, P, WAG, are thanked for allowing access to study their *Cola* specimens and/or sending loans of specimens to K, especially Paul Musili, Kennedy Matheka, Geoffrey Mwachala at EA and Emile Kami at IEC.

The author declares that he has no conflict of interest.

